# A deep-learning tool for species-agnostic integration of cancer cell states

**DOI:** 10.1101/2024.12.20.629285

**Authors:** Jonathan Rub, Jason E Chan, Carleigh Sussman, William D. Tap, Samuel Singer, Tuomas Tammela, Doron Betel

## Abstract

Genetically engineered mouse models (GEMM) of cancer are a useful tool for exploring the development and biological composition of human tumors and, when combined with single-cell RNA-sequencing (scRNA-seq), provide a transcriptomic snapshot of cancer data to explore heterogeneity of cell states in an immunocompetent context. However, cross-species comparison often suffers from biological batch effect and inherent differences between mice and humans decreases the signal of biological insights that can be gleaned from these models. Here, we develop scVital, a computational tool that uses a variational autoencoder and discriminator to embed scRNA-seq data into a species-agnostic *latent space* to overcome batch effect and identify cell states shared between species. We introduce the latent space similarity (LSS) score, a new metric designed to evaluate batch correction accuracy by leveraging pre-labeled clusters for scoring instead of the current method of creating new clusters. Using this new metric, we demonstrate scVital performs comparably well relative to other deep learning algorithms and rapidly integrates scRNA-seq data of normal tissues across species with high fidelity. When applying scVital to pancreatic ductal adenocarcinoma or lung adenocarcinoma data from GEMMs and primary patient samples, scVital accurately aligns biologically similar cell states. In undifferentiated pleomorphic sarcoma, a test case with no *a priori* knowledge of cell state concordance between mouse and human, scVital identifies a previously unknown cell state that persists after chemotherapy and is shared by a GEMM and human patient-derived xenografts. These findings establish the utility of scVital in identifying conserved cell states across species to enhance the translational capabilities of mouse models.

## Introduction

Critical insights into human biology and cancer have emerged from the use of model organisms. Genetically engineered mouse models (GEMMs) or transplantable mouse tumor models underpin virtually every clinical success story in oncology. However, mouse cancer models harbor fundamental differences to humans, including species-specific malignant differentiation states. These differences have limited accurate prediction of pathophysiology and treatment outcomes of cancer in human patients. It is estimated that only about one third of the research performed in animal models proceed to clinical trials in humans. Of those trials, less than 10% are successful in Phase I [1], and of those that pass Phase I, less than one in ten are clinically approved by the FDA—a dismally low fraction [2].

The use of GEMMs in the research of rare cancers is particularly important. GEMMs provide an avenue to investigate these diseases in a controlled and reproducible fashion, thereby abrogating one of the main issues of rare cancer research – the availability of sample material to study. As such, an unmet need exists to improve the predictive value of pre-clinical mouse models of cancer. Solid tumors are composed of functionally and molecularly distinct cancer cell subpopulations. The clinical relevance of this intratumoral heterogeneity manifests in the distinct capacities of cancer cell states for growth, progression, and treatment resistance. Thus, understanding the extent to which mouse models recapitulate the cell state heterogeneity of human tumors is of critical importance. Furthermore, cell states that are conserved across species are likely to be biologically and functionally important, motivating their identification.

Improving the accuracy of pre-clinical models is of particular importance in the field of sarcoma, which are a set of rare cancers affecting the soft tissue and bone. These cancers make up only one percent of new cancer diagnosis each year in the United States [3]. Sarcomas are a very diverse group of cancer and many subtypes, such as undifferentiated pleomorphic sarcoma (UPS), harbor a dismal prognosis [4]. The incidence of UPS in the United States is about 1 in 200,000 [5]. Due to its rarity and severity, interrogation of sarcoma biology is often difficult. Clinical trials often require coordination of multiple institutions and can take many years. In these cases, GEMMs are especially useful to generate data to understand rare cancer biology in order to develop appropriate treatment strategies, yet it is unclear how well these models recapitulate the biology and cell state heterogeneity of human UPS [6].

Computational modeling approaches, such as deep learning, hold considerable potential for increasing the predictive power of animal models of human cancer [7] [8]. Single cell RNA-sequencing (scRNAseq) provides an unbiased snapshot of the cellular composition and associated single-cell gene expression programs in tumors. This sensitive method has been used to elucidate cancer cell states and gene programs across patients [9] [10]. However, computational methods specifically designed to discern similarities and differences across mouse models and human cancer have not been developed [7] [8].

Previous approaches for cross-species comparison of scRNA-seq are typically akin to scRNA-seq batch correction approaches where differences between datasets are dominated by technical variability from experiments. A common approach for cross-species scRNA-seq comparison is to apply widely used batch correction algorithms like BBKNN [11], Harmony [12], scVI [13] or scDREAMER [14]. However, these algorithms do not specifically address cross-species integration of homologous samples where a subset of the cells may be indistinctly different between species while other cell types are highly homologous between species. Another common practice in cross-species cancer studies is to analyze each species dataset separately, followed by comparison of marker genes or specific gene signatures [15]. These current approaches are limited to genes that have one-to-one homology between two species. Thus, they likely fail to identify functionally or biologically similar species-specific cell states that are driven by species-specific genes.

We hypothesized that computational methods can help decipher transcriptomic differences between mouse cancer models and human cancers to identify core features of cancer conserved in human tumors. Identifying similar cancer cell states across species will increase the clinical relevance of studies utilizing mouse models. Here, we introduce a new approach for cross-species integration without a limitation to species-specific genes based on **s**ingle-**c**ell **V**ariational autoencoder **i**ntegration **t**hrough species-**a**gnostic **l**atent space (scVital) to bridge this technology gap.

## Results

### ScVital model

ScVital uses a variational conditional autoencoder (VAE) [16] coupled with an adversarially trained discriminator to embed scRNA-seq data from different species into a common latent space (Fig. 1A). VAEs take high-dimensional input data and embed the data into a smaller, generalizable latent dimension that is then used to reconstruct the original high-dimensional data. In the context of scRNA-seq, the VAE takes as input cells represented as vectors of high-dimensional gene expression data, which is computationally inefficient to analyze directly, and maps the data into a smaller, low-dimensional latent space representation that captures the salient features of the cell state [17]. To exclude the species-specific signal from the latent space representation, scVital simultaneously trains a discriminator network to predict the species from which the latent space originated, a concept adopted from generative adversarial networks (GANs) [18]. The output from the discriminator is then incorporated into the VAE training function such that the resulting latent space representation preserves cellular identity but excludes species signatures (see Methods section for details).

**Figure 1:**
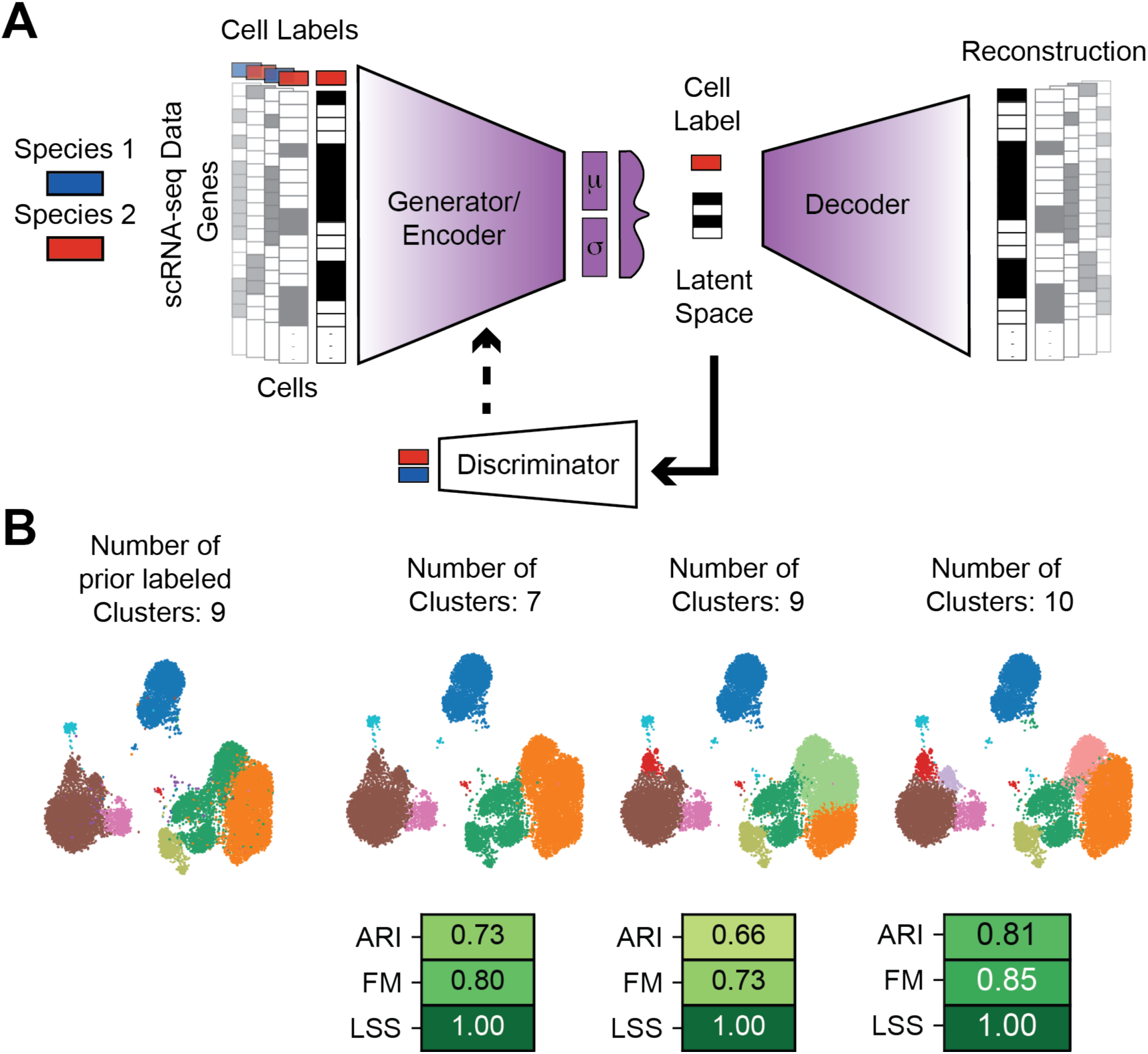
ScVital architecture and latent space similarity (LSS). **A.** ScVital architecture schematic. ScVital is made up of a variational autoencoder and discriminator. ScVital takes batch labeled scRNA-seq data as input and outputs a batch agnostic latent space. A variational autoencoder (VAE) is trained by minimizing the mean squared error of the input scRNA-seq data and reconstructed scRNA-seq data, minimizing the KL divergence of the latent space, and maximizing the discriminator cross-entropy loss. The discriminator is trained by minimizing the cross-entropy loss of the predicted batch labels and true batch labels. The resulting latent space can be used as a batch agnostic embedding of the data for downstream analysis. The full architecture is further explained in the *Methods* section. **B.** Integration metrics are sensitive to the number of clusters. UMAP visualization of scRNA-seq data with different number of clusters, generated by different clustering parameters, showing the variability of ARI and FM scores to the number of clusters and inadequacy of these metrics as measures of integration. The far left UMAP is the pre-integration labeled clusters (9 clusters). ARI and FM scores change drastically with the post-integration clustering results while the LSS scores is consistent and independent of ad-hoc clustering.

To test the performance of scVital, we compared it against several current batch correction algorithms: BBKNN [11], Harmony [12], scVI [13], and scDREAMER [14] using multiple datasets of varying complexity. Both scVI and scDREAMER are based on a similar VAE approach as implemented in scVital although they are primarily designed for data integration. Often two-dimensional visualizations, like uniform manifold projection (UMAP), aid in scRNA-seq analysis; However, this two-dimensional projection is a warped view of the high-dimensional gene expression space meant to be used as a visualization tool, not as a metric for analysis of similarity or integration. Therefore, we used more robust metrics to score the accuracy of scVital integration.

### Model evaluation metrics

A common approach to evaluate integration of single cell datasets is to use adjusted rand index (ARI) [19] [20] and Folks-Mallow (FM) scores [21] to determine cell type conservation [22] [23]. ARI and FM measure the concurrence between two clustering results by calculating the ratio of agreed versus discordant clusters. These measures score the integration results by comparing the prior cell labels to a set of new cell labels defined after integration. In this case, they measure the similarity between the true, previously labeled cells and the predicted, to the newly labeled cells from the output of the integration algorithms. The limitation of ARI and FM is that they require labeling of the post-integration clusters and therefore they are highly sensitive to the number of clusters. Clustering single cell data is highly variable, imprecise, and often heuristic, depending on the choice of algorithm and parameters [24] [25]. To avoid this dependency on heuristic clustering for evaluating and comparing integration, we developed a latent space similarity score (LSS) as a metric that can measure the integration using only the known, true cell labels without the need for new cell labels. (Fig. 1B). This approach computes the pairwise cosine similarity between the original (non-integrated) cell labels in the integrated latent space (see Methods). It then calculates the area under the curve of the F1 score (AUC-F1, the harmonic mean of the precision and recall) to determine the accuracy of integration in correctly pairing common cell labels in the latent space. Hence, the integration score is independent of new clustering of cells after integration and simply evaluates whether cells with common labels are closer in the integrated latent space relative to other cells.

### Integration of mouse and human normal tissue data

We find scVital is comparable to other gold standard scRNA-seq integration algorithms, such as Harmony and scVI, when integrating common batch correction datasets, including peripheral blood monocytic cell (PBMC) data [11] sequenced from 5’ and 3’ ends of transcripts (Supplementary Fig. S1). ScVital also works comparably well for integrating mouse and human scRNA-seq data from multiple healthy tissues including from muscle [26] [27] (Fig. 2), lung [28] (Supplementary Fig. S2), pancreas [29] (Supplementary Fig. S4), liver [30] [31] (Supplementary Fig. S3), and bladder [15] (Supplementary Fig. S5).

**Figure 2:**
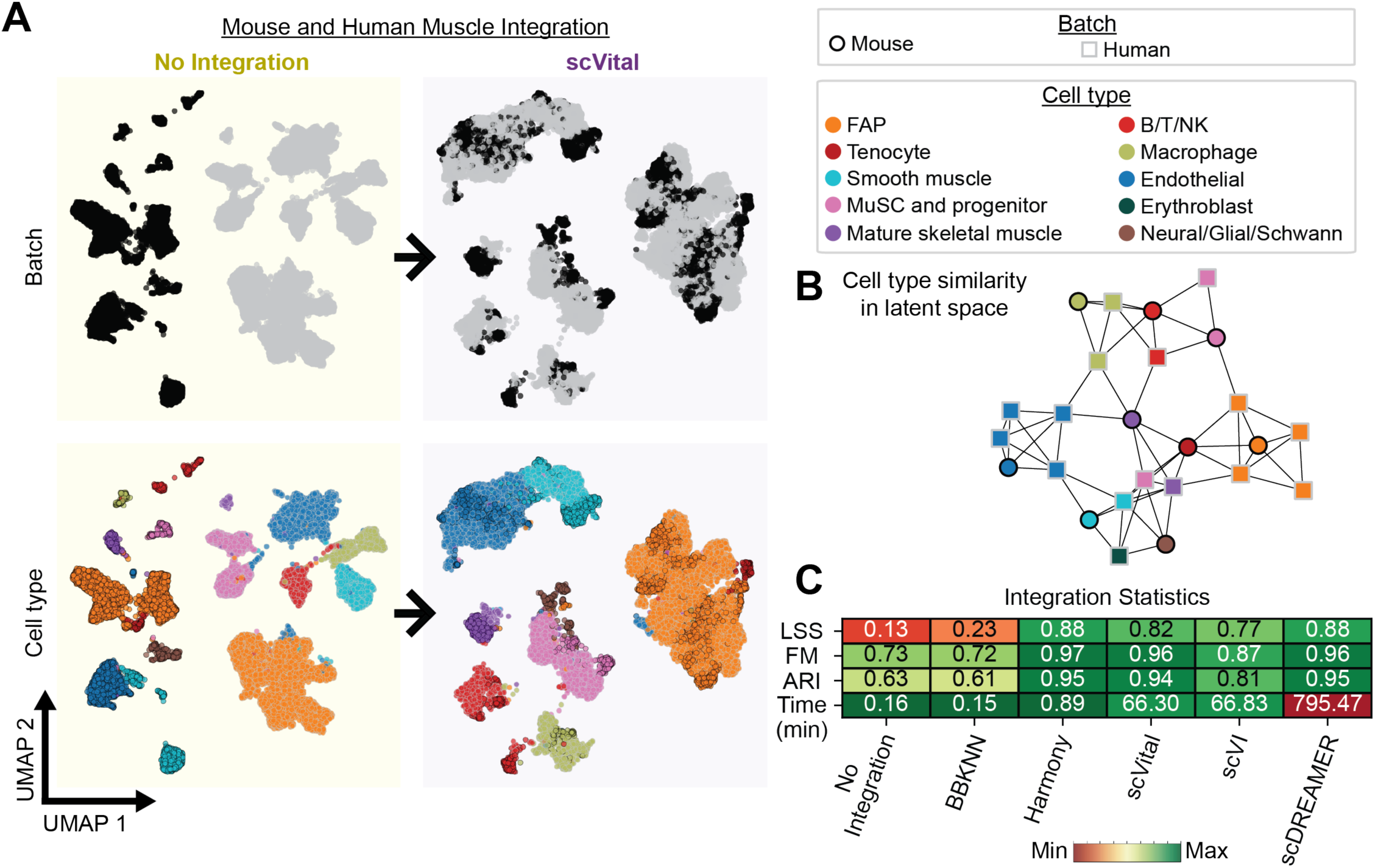
Integration of mouse and human muscle data. **A.** UMAP visualizations of mouse and human muscle data with no integration in yellow (left) and following scVital integration in purple (right) colored by species (top) and cell type (bottom). Cell point outlines represent the species (black for mouse and gray for human) and internal color represents the cell type (ex. smooth muscle in light blue). With no integration there is a clear separation of mouse and human muscle data and no overlap of homologous cell types. After integration with scVital similar cell types are overlapping as shown by the overlapping color tones in the UMAP visualization. Fibro-adipogenic progenitor cell (FAP), B cell/T cell/NK cell (B/T/NK), and muscle stem cell and progenitor cell (MuSC and progenitor). **B.** Graph visualization of LSS. Each node represents the average latent space of a cell type after scVital integration where the outline color is the batch and internal color is the true label as determined by the input dataset. The distances between nodes represent the cosine similarities of the cell type’s average latent space compared to other cell types. Connected same-colored nodes with black and gray outlines indicates integration of species and conservation of cell type. For example, the blue nodes denoting endothelial cells are most similar to each other with both square, black-outlined human nodes connected to circular, gray-outlined mouse nodes. **C.** Integration scores, LSS, FM, ARI and runtime (in minutes) of mouse and human muscle with no integrations and integrated using, BBKNN, Harmony, scVital, scVI, and scDREAMER. ScVital performs comparably well to the other gold standard integration algorithms achieving similar ARI, FM, and LSS scores.

UMAP representation of mouse and human normal muscle scRNA-seq data without integration illustrates the strong species effects on data integration, whereas following scVital integration the species effect is removed while the cell type information is maintained (Fig. 2A). A graph visualization of the LSS (i.e. the cell type similarity values) further reinforces the UMAP view and demonstrates closer similarities among homologous mouse and human cell types than other cell types (Fig. 2B). ScVital metrics of integration are comparable to other integration algorithms (Fig. 2C) and its runtime performance is significantly shorter than the other deep learning algorithm scDREAMER while displaying better or comparable integration metrics across all mouse and human normal datasets (Supplementary Figs. S1C, S2C, S3C, S4C, and S5C).

### Integration of mouse and human cancer data

Malignant cell states are typically more challenging to define and distinguish from each other compared to non-malignant cell types given they arise via dynamic and constantly occurring transdifferentiation of cells within the same lineage. GEMMs and allograft models provide a means to model cancer cell states in an isogenic system in vivo. Yet, the extent to which these models approximate human tumors remains unclear due not only to cross-species biological differences but also interpatient heterogeneity. A primary motivation for this work is to identify the core features of cancer GEMMs and transplantable models that are conserved in human tumors by identifying similar cancer cell states across species. To determine how scVital performs when integrating malignant cell states, we performed cross-species integration of primary human tumors and GEMM tumors of pancreatic ductal adenocarcinoma (PDAC), lung adenocarcinoma (LUAD), and undifferentiated pleomorphic sarcoma (UPS).

PDAC is composed of 2 dominant cell states, one with classical epithelial features and the other with a basal identity [32] [33] [34]. PDAC GEMMs show high concordance between the human classical and basal states, but in addition harbor a mesenchymal state that is not commonly found in human primary PDAC [35] [32]. We integrated scRNA-seq data from two mouse GEMM studies [36] [35] and from 24 human [37] primary tumors patient samples to measure the accuracy of scVital and its ability to identify common cancer cell states (Fig. 3). Before integration, the PDAC samples exhibited a strong cross-species effect with significant interpatient differences (Fig. 3A). Integration with scVital yielded species-agnostic alignment of classical and basal states, whereas the mouse mesenchymal cell state separated from the human cell states (Fig. 3B). ScVital integrates multiple PDAC datasets with a comparable LSS score of 0.66 to Harmony (0.62), and scVI (0.60). We note that scDREAMER had the highest LSS score (0.91) but suffered from a runtime of 1,311 min, 1,171 min slower than scVital (Fig. 3C). Further, after integration scVital pointed to similarity of the basal cell state to the mesenchymal cell state, which has been suggested by previous scRNA-seq studies [32]. Taken together, these results establish the ability of scVital to identify and align previously known biologically conserved cell states across species in PDAC.

**Figure 3:**
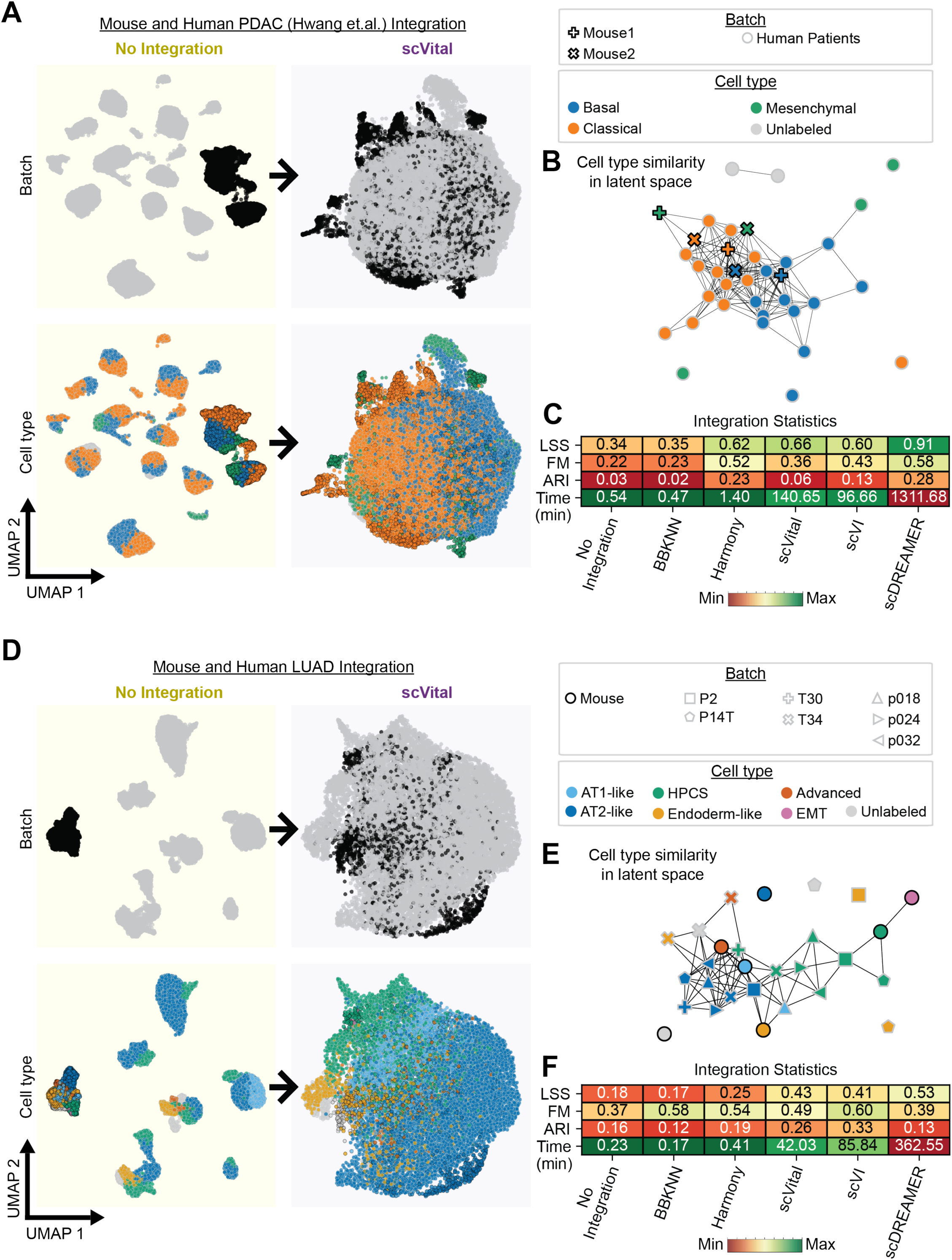
Integration of mouse and human LUAD and PDAC data. **A.** UMAP visualizations of GEMM and human patient PDAC data with no integration (left) and after scVital integration(right) colored by species (top) and cell type (bottom). Without integration the data segregates by sample source (human, mouse). Integration with scVital identified common classical, basal cell states, and mouse-specific mesenchymal PDAC cell states. **B.** Graph representation of LSS scores of the PDAC data shows the separation of classical, basal, and mesenchymal cell states. Node shapes denote sample origin and colors according to cell type label. The graph shows high similarity of the basal cell state between human patients and mouse model and separation of the basal and classical cell states. **C. & F.** ScVital has comparable integration metrics to scDREAMER and scVI with significant faster runtime. **D.** Similar to **A,** UMAP views of LUAD from 7 human patients from 3 datasets and a LUAD GEMM showing scVital integration identifies common of AT2-like and HPCS cell states across human patients and GEMM. **E.** Similar to **B,** LSS graph of the LUAD data shows the separation of LUAD cell states depicts HPCS (green) similarity between GEMM and human.

We next applied scVital to LUAD scRNA-seq data, a disease characterized by high intra-tumoral and inter-patient heterogeneity [38]. First, we aggregated LUAD data from a GEMM study [39] and eight patient samples from three different human LUAD studies [40], [41], [42]. Without integration, the LUAD samples showed minimal overlap and cells clustered according to species and based on patient. (Fig. 3D). Integration of the samples with scVital identified overlap between a cell state with similarity to alveolar type 2 cells (AT2-like state) and a high plasticity cell state (HPCS) across patients and cross-species [38] (Fig. 3D, E). A similar analysis of integrating mouse and human healthy lung data showed high concordance of mouse and human AT2 cells (Supplementary Fig. 2). Again, the runtime of scVital was faster compared to scVI and scDREAMER (42 min versus 85 min and 362 min, respectively) with a comparable LSS score (0.43 versus 0.41 and 0.53, respectively) (Fig. 3F).

### Integration of disease and normal tissue data

We tested the ability of scVital to integrate and discriminate cell states and cell types between healthy tissue and different diseased cell states. Such a comparison can identify commonalities and differences between normal cells, regenerative processes, and disease states. We integrated scRNA-seq data from mouse non-tumor lung [28], alveolar injury [43], and LUAD samples [39] (Supplementary Fig. S6). We found significant overlap between AT2 cells from healthy lung, AT2 cells from injured lung, and AT2-like cells from LUAD samples. Interestingly, the integration uncovered high similarity between the mouse LUAD HPCS and a damage associated transient progenitor (DATP) cell state, a transitory cell state associated with lung injury absent in quiescent, healthy lung (Supplementary Fig. S6). As expected, ciliated and neuroendocrine cells did not match any of the alveolar injury-associated or LUAD cell types.

We performed an analogous comparison of human LUAD and healthy human lung tissue (Supplementary Fig. S7). Similar to the mouse, this analysis identified high similarity between the non-malignant alveolar cell populations and the malignant AT2-like cell state, while the human basal cells remained separate from LUAD cell states. The malignant human HPCS cells show similarity, but not integration, with other non-malignant cell states, possibly because the HPCS resembles the DATP that is associated with injury—a cell state that was absent in our healthy human lung dataset [43]. Similar to the mouse results, the overlap of non-malignant AT2 cells and the malignant AT2-like cells in mouse and in human can be attributed to the shared lineage of LUAD to AT2 cells, which are the predominate LUAD cell-of-origin [44].

### Integration of mouse and human UPS data

To explore the utility of scVital in a case where there is minimal prior knowledge about the cellular composition, we used scVital to integrate malignant states in a GEMM of undifferentiated pleomorphic sarcoma (UPS) with patient-derived xenograft (PDX) models. UPS is an aggressive, rare soft tissue sarcoma with complex inter- and intratumoral heterogeneity. Recent studies identified these tumors to contain complex genetics, with only TP53 as a commonly mutated gene across patients [45] [46] [47]. The complex genetics and high inter-tumoral heterogeneity, in combination with the rarity of the disease, make it imperative to use mouse models to gain an in-depth understanding of the biology of UPS. At present, the most widely used UPS GEMM relies on Cre recombinase-mediated activation of ***K****ras^lox-stop-lox-G^*^12*D/*+^ and *Tr* ***p****53^flox/flox^* alleles (**KP** model) [6]. This animal model results in tumors that are histologically similar to UPS in human [6]. Though the *Kras^G^*^12*D*^ mutation is rarely present in human UPS tumors [45] [46] [47], approximately 86.5% of UPS cases show hyperactive MAPK signaling, which is associated with a poor prognosis [48].

To interrogate features of cell state dynamics conserved across mouse and man, we treated KP GEMM UPS tumors and UPS PDXs with short- or long-term doxorubicin, a common chemotherapy backbone used to treat advanced-stage soft tissue sarcomas [49], followed by scRNA-seq analysis (Fig. 4A). Without integration tumors from the three UPS models (KP GEMM and two PDXs) were distinct and did not appear to share cellular states (Fig. 4B). Notably, only integration with either scVital or scDREAMER demonstrated overlapping cell populations across all three datasets, whereas other integration tools failed to fully integrate the data sets (Fig. 4C).

**Figure 4:**
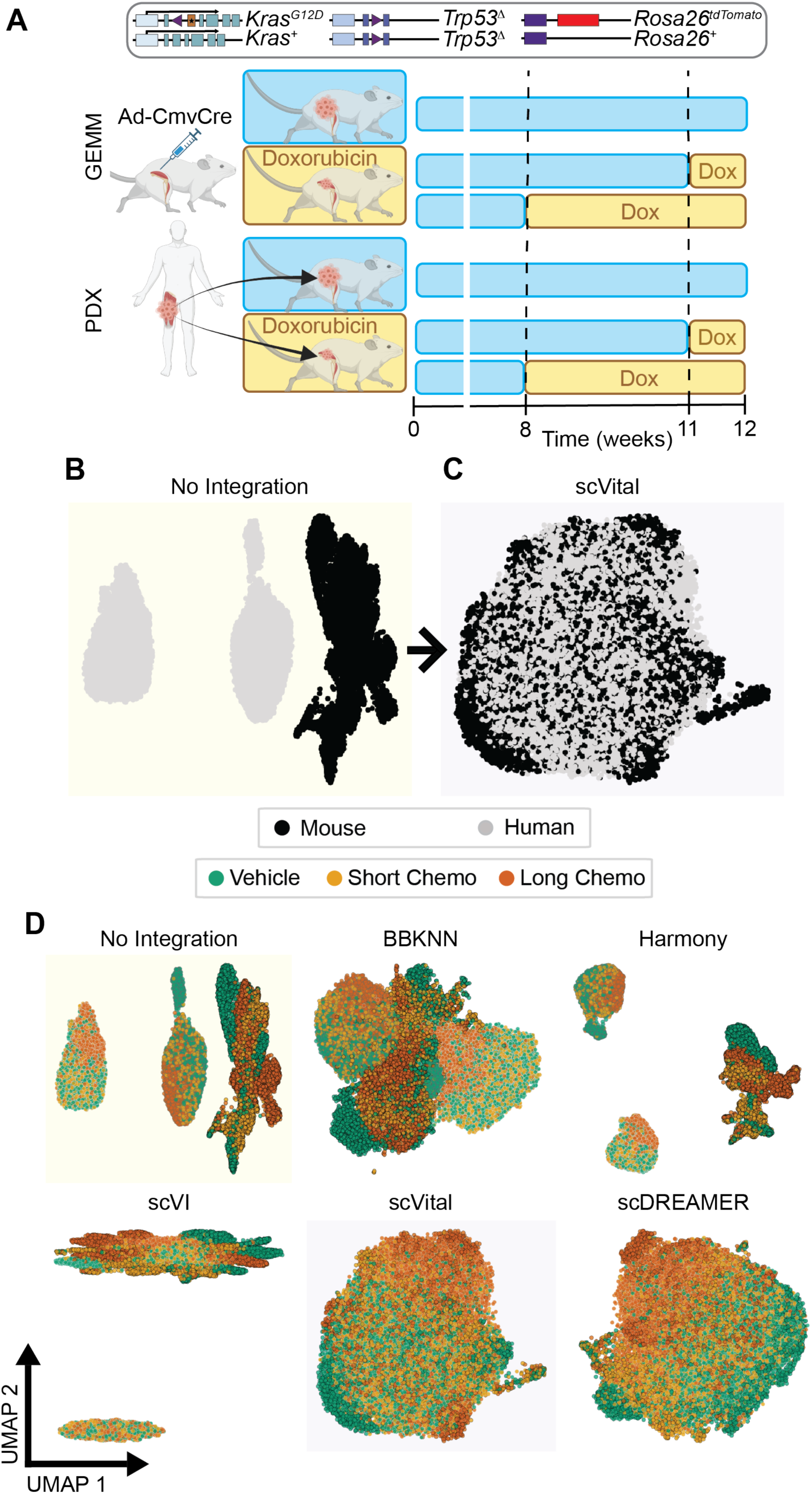
Integration of GEMM and PDX of UPS. **A.** Treatment experiment outline. KP GEMM mouse model of UPS and PDX of UPS were treated with vehicle, short-term doxorubicin (one week), or long-term doxorubicin for scRNA-seq analysis. (see *Methods* for more details). **B.** With no integration there is no overlap between the two PDXs and the GEMM. Each sample clusters on its own and no similarities are identifiable. After integration with scVital there is overlap of the PDXs and GEMM datasets. **C.** Integration results of UPS GEMM and PDX samples by different algorithms demonstrates that scVital and scDREAMER are the only approaches that are able to integrate the two PDX and GEMM and find overlapping cell states, albeit with significant difference in runtime performance. (scVital: 31min, scDREAMER: 651min)

We identified one common cell state dominated by cells from the long-term treatment group across both species (Fig. 5A, B). This cluster is enriched with a hallmark hypoxic gene signature [34]. Investigating this overlapping population further we identified the conserved hypoxia marker *SLC2A1/Slc2a1* [50] (5C). Staining of KP or UPS PDX tumors revealed a similar significant increase in SLC2A1 protein expression in doxorubicin-treated samples compared to vehicle control (Fig. 5E, F). The quantification of the staining further reinforced the findings, demonstrating increased expression of the hypoxia markers in the extended treatment for both the GEMM and PDX (Fig. 5G, H). Notably, the commonly used practice to analyze the three UPS tumor datasets separately and then use the intersection of marker genes that characterize each treatment group would have not identified a similar hypoxic signature, including the hallmark *SLC2A1/Slc2a1* marker that defined that overlap (Supplementary Fig. S8). These results highlight the power of scVital in identifying biologically and functionally similar cancer cell states in tumors.

**Figure 5:**
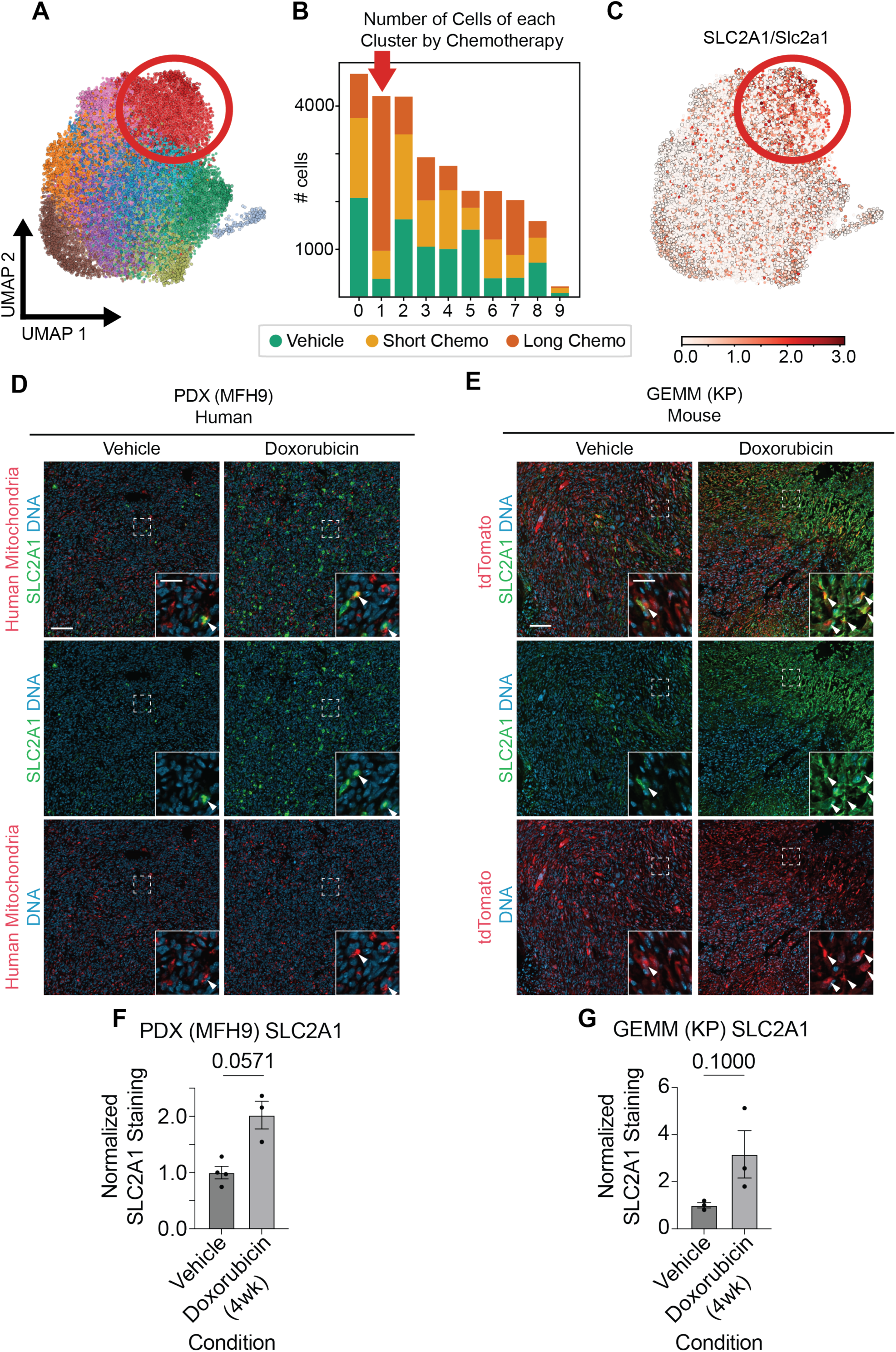
Identification of hypoxia signature in treatment resistant UPS cells. **A. & B.** Longterm doxorubicin resistant cells were identified following scVital integration. Clustering analysis identified cluster 1 (circled in red) (B) enriched in long-term treated samples. **C.** UMAP of UPS data visualizing *SLC2A1/Slc2a1* expression. *SLC2A1/Slc2a1*, a common hypoxia marker, is expressed mostly in cluster 1 (circled in red on the UMAP). **D.** Histological staining of *SLC2A1* in human PDX. Outer scale bar: 100 *µm*, inset scale bar: 25 *µm*. UPS cells, identified with human mitochondria staining, treated with vehicle have little to no *SLC2A1* expression. After treatment with doxorubicin there is an increase in *SLC2A1*. The cells marked by arrows in the inset show an example of a human UPS cell expressing *SLC2A1* after doxorubicin treatment. **E.** Histological staining of *SLC2A1* in mouse GEMM. Outer scale bar: 100 *µm*, inset scale bar: 25*µm*. UPS cells, identified with *tdTomato* staining, treated with vehicle have little *SLC2A1* expression. The inset identifies one cell with co-expression of the markers. After treatment with doxorubicin there is an increase in staining of *SLC2A1*. The cells marked by arrows in the inset show an example of a mouse UPS cell expressing *SLC2A1* after doxorubicin treatment. **F & G.** Quantification of the staining of SLC2A1 expression in the PDX and GEMM of UPS. There is an increase in SLC2A1 expression after doxorubicin treatment in both the human PDX and the mouse GEMM (Mann-Whitney U; n >= 3 mice each group).

## Discussion

Mouse models of cancer are pivotal in understanding human disease. At present, it is unclear how well current methods identify overlapping cell states in mouse models and human cancer. In addition, current methods do not perform integration with species-specific genes potentially overlooking biologically relevant species-specific cell states. Here, we show the utility and performance of scVital by integrating multiple mouse and human scRNA-seq datasets. Our method, scVital, uses an autoencoder and discriminator to embed scRNA-seq data of homologous and species-specific genes into a shared latent space. We tested the accuracy of scVital by performing integration of healthy mouse and human tissue including muscle, liver, lung, pancreas, and bladder where scVital was able to correctly integrate common cell types. We then performed integration of GEMM and human cancer data from PDAC, LUAD, and UPS tumors. ScVital was able to accurately and quickly integrate the data and find overlapping cancer cell states.

Specifically, when applying scVital to GEMM and human PDX of UPS we identified a cell state enriched for a hypoxia signature that is shared in the mouse model and human PDXs. Only one other integration method, scDREAMER, was able to find a similar overlap however, scVital runtime is more than 10 times faster. Hypoxia has previously been associated with chemoresistance to doxorubicin [51] [52] and several studies have suggested that hypoxia is associated with poor prognosis in UPS [53] [54] [55]. Identification of this cell state and its conservation in GEMMs enables functional studies on this cell state, the results of which can be extrapolated to human disease. This is particularly significant in rare cancers, such as UPS, where clinical trials are difficult to execute due to the rarity of cases and high inter-patient heterogeneity of the disease.

Both Gavish et al. [10] and Barkley et al. [9] performed a pan-cancer analysis and found conserved cell states across different tumors in human patients, referred to as meta-programs and modules, respectively. By using scVital to integrate tumor data across species we are able to draw similar conclusions and find overlapping cell states across species. For example, Gavish et al. identified a PDAC classical meta-program that is present across multiple tumor types. When integrating PDAC data, we similarly identified conservation of the classical cell state across species. In addition, the high-plasticity cell state (HPCS) in LUAD GEMM also exists in human LUAD [38]. Using scVital, we corroborated these findings by integrating GEMMs and multiple patients with PDAC or LUAD.

When comparing integration of scVital with other gold standard scRNA-seq batch correction methods, we find that the other methods are also able to accurately integrate normal mouse and human tissue. However, the common batch correction algorithms performance decline with integration of malignant cell states which are challenging to discern from each other since they arise via cells from the same lineage that are constantly evolving and changing. Harmony is a very fast algorithm that is able to integrate scRNA-seq with high accuracy and speed in most cases [22]. However, Harmony did not accurately integrate cell states when analyzing GEMM and human PDX UPS data, indicating it is limited when integrating highly complex data.

We introduce latent space similarity (LSS), a new metric to accurately determine validity of integration when cells are pre-labeled. This metric is a cluster-agnostic method of scoring integration. When used together with ARI and FM, LSS is a powerful tool for determining integration accuracy. For example, high LSS and low ARI and FM signifies a high capability for integration, as determined by the latent space, but the post-integration clustering does not capture the new cell labels correctly which leads to low ARI and FM scores. High LSS, ARI, and FM scores indicate accurate integration and accurate post-integration clustering of the cells. Low LSS values may indicate that the data sets contain distinct, non-homologous cell states suggesting the cell type pairings are wrong, or that the original cell labels are incorrect. In the latter case, LSS score can also be used as quality control measure of scRNA cellular annotation. This metric alongside existing integration measures is helpful to determine accuracy of integration and visualize the similarity of cell types. A limitation of LSS is that the data must be pre-labeled and the labels need to be paired between data sets prior to scoring. We note that this is a limitation of all scoring metrics as ARI and FM also require known labels and pairings to perform scoring.

Future iterations of the architecture of scVital could be made in effort to improve integration and down-stream analysis by incorporating additional confounding factors. For example, inter-patient heterogeneity is also a source of batch effect when integrating human data. A potential future development of scVital could be to add another discriminator network so that one discriminator could remove the patient batch effect and the other would remove the species batch effect. This addition could improve scVital integration when multiple human patients are being integrated. Another avenue of exploration of scVital is to use the reconstructed output data as an imputed data to overcome gene dropout that often occurs in scRNA-seq data [56]. This imputed gene expression data can then be used for further downstream analysis, including differential gene expression. Lastly, scVital could be expanded to perform cell clustering. By adding an output clustering layer to the latent space, the input data would be clustered based on similarity in the latent space. This clustering would help identify cell states without the need for Leiden clustering after training.

The inherent differences between mouse and human cancers can limit the utility of mouse models. Here we introduce scVital as an integration tool that can help bridge those differences to identify conserved human and mouse cell states that are likely to be therapeutically important. This increase of knowledge transfer can help us understand cancer progression in humans with mouse model experiments, which have distinct advantages, such as tracing and eventual ablation of distinct cell states [57], perturbation of genes of interest [58], and temporal analyses using time points. ScVital is likely to improve the predictive power of *in vivo* animal models of cancer for translational studies and clinical trials.

## Materials and Methods

### scVital Architecture

ScVital is variational autoencoder (VAE) neural network model made up of three neural networks: an encoder, a decoder, and a discriminator (Figure 1). The input to the model is the vector of log-normalized highly variable cellular gene expression values {*x*_1_*, x*_2_*, x*_3_*, …, x_G_*} with an additional value *x_G_*_+1_ representing the sample source (i.e. species, sample, or batch). The encoder and decoder are symmetric with two hidden layers. The encoder and decoder layers are fully connected to normalization layers [59] and a ReLU activation function. The number of nodes for the internal layers is fully customizable with default value of 1024 and then 128. The latent space has a default size of 12 nodes. The variational autoencoder outputs 24 (12*2) nodes, corresponding to the mean and the natural log of the variance. The decoder takes in the latent space and conditional nodes as input and outputs a reconstruction of the original dimensional scRNA-seq data. The default overall architecture, with 2000 input genes integrating two batches is: 2002 -> 1024 -> 128 -> 24 -> [latent space of 12] -> 14 -> 128 -> 1024 -> 2000.

The discriminator takes the latent space values as input and outputs probabilities for the latent space to have originated from each batch. The cross-entropy between these predicted probabilities and the latent space’s true origin is used to calculate the loss value and train the discriminator. The discriminator has one fully connected hidden layer with a ReLU activation function. When there are two batches the default architecture for the discriminator is 12 – 6 – 2. This architecture is flexible to any number of batches, the output dimension increases for number of batches.

Both the VAE and discriminator are trained using adaptive moment estimation [60] with weight decay (AdamW) optimizer coupled with cosine annealing with warm restarts learning rate scheduler [61]. The AdamW optimizer implements weight decay for the Adam optimizer more faithfully and when paired with cosine annealing. The cosine annealing with warm restarts learning rate scheduler cycles the learning rate of the optimizer from 1*e^−^*^3^ to close to 0 then back up to 1*e^−^*^3^ after a certain number of epochs, in this case five epochs. This optimizer helps train a network with a wider set of values that provide optimal results. This method allows the network to avoid converging to a local minimum during training and prevents converging to a local minimum during training [62].

The discriminator component uses the latent space to generate batch probabilities. The autoencoder and discriminator are adversarially trained in a min-max approach where the discriminator is trained to accurately distinguish the latent space between the different batches (by minimizing the cross-entropy loss between the true labels and the predicted labels) and the autoencoder creates a latent space that is indistinguishable between batches by minimizing the loss between the original and reconstructed data, minimize the Kullback-Leibler (KL) divergence, and maximize the discriminator loss. The conditional node in the autoencoder input and the discriminator ensures the latent space is species-agnostic where a latent space represent embedding of common cell states across species. Downstream scRNA-seq analysis can be performed on the latent space from scVital.

### Loss functions

We denote expression levels of gene *g* in cell *x* as *x*_{*g∊G*}_, and accordingly denote the entire scRNA-seq dataset of all cells and genes in the dataset as **X***_G_*, where *G* is total number of genes and *N* is the total number of cells.

ScVital is trained using a loss function (e.q. 1) made up of three weighted components with default weight values determined by coarse grid search using several benchmark datasets.

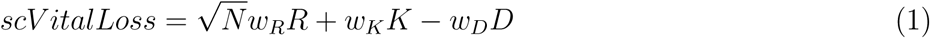

1. The first component is a reconstruction loss (*R*, eq. 2) which is calculated using the mean squared error between the original and the reconstructed scRNA-seq data from the autoencoder. Gene homology between batches are identified using the mouse genome informatics resource [63], and the names are then harmonized for downstream analysis. A novel aspect of the training scVital is that reconstruction loss uses the entire species gene set, including the non-homologous, when calculating the reconstruction loss. This means that human specific genes are ignored when calculating the reconstruction loss of mouse cells. The coefficient is also augmented by multiplying the value by the square root of the input data size. The weight coefficient *w_R_* controls the contribution of cell type identification to the overall cost; where high values of this coefficient will result in fragmented cell clusters with limited species overlap whereas low values result in integrating unrelated cells. The default value is 2, defined empirically providing optimal results in most test cases and is scaled by the square root of the size of the input data.

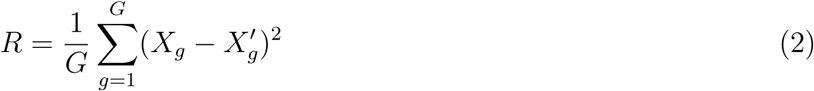

2. The second loss function is KL divergence loss (*K*, eq. 3), which is calculated by determining the divergence of the latent space of size *L* with the normal distribution 𝒩 (*µ* = 0*, σ*^2^ = 1) where *σ_l_* and *µ_l_* are output from the last layer of the encoder from the autoencoder. This component insures a regularized network to avoid overfitting. The weight coefficient of *w_K_* controls the tradeoff between lack of generalization (i.e. overfitting the model) versus over-generalization leading to random prediction and loss of predictive power. We use a cyclical annealing coefficient during training where initially the *w_K_* = 0 and is gradually increased over a customizable number of epochs (default value of 5) then decreased again. This allows the autoencoder to have a higher weight on the reconstruction loss then gradually increases the regularization by increasing *K* coefficient [64].

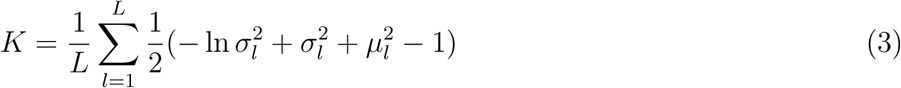

3. The third loss function is the discriminator loss (*D*, eq. 4), which is used to predict the species (or batch) *b* of the cells. It is calculated using cross entropy loss between the batch label and the discriminator output of the species probability derived from the latent space. The negative coefficient *w_D_*determines the balance between removing the species origin while preserving the cell type identity when training the autoencoder.

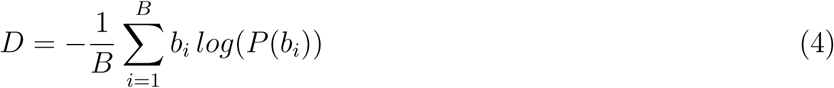

Default values for the three *w_i_* weights were determined by coarse grid search using several benchmark datasets. Overall scVital performance is most sensitive to *w_R_* contribution.

### Latent space similarity

Most metrics to determine accuracy of integration rely on preservation and matching of cluster identity pre and post integration. A common measure is the Adjusted Rand Index defined as 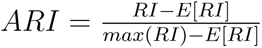 where 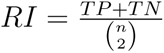 and *TP* is number of cases where pre integration cluster labels match post integration labels and conversely *TN* are the number of correctly un-matched labels and *n* is the number of clusters. This measure is heavily dependent on the sample post integration clusters and accuracy of the cell labels, both are unresolved challenges in single cell analysis. To avoid this dependency on clustering of the cells we developed an integration metric that determines the accuracy of integration based on the similarity in the shared latent space.

Latent Space Similarity (LSS) is the average latent space cosine distance, 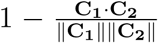, between all the cells of type *C* from each datasets (*C_i∊_*_1,2_). Intuitively, cosine distances between homologous cell types (e.g. between mouse fibroblast and human fibroblasts) are expected to be lower than between any other cell types (even within the same organism). The cell type cosine distances are used to calculate the AUC-F1 performance of the integration using predetermined pairs of cell types to determine the accuracy of integration of the cell types in the latent space. AUC-F1 is the harmonic mean of the precision and recall, where the precision is the number of true positives (TP) divided by the number of true positives and false positives (FP) and the recall is the number of true positives divided by the number of true positives and the false negatives (FN). 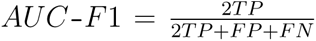. F1 values can range between 0 and 1, where values close to 1 signify cell type pairs had the lowest cosine distance and accurate integration across species whereas values close to 0 signify random cell type pairing and poor integration. AUC-F1 metric evaluates if the integration was able to overcome the species effect and determines the accuracy of cell types integrating with the corresponding similar cell type across species instead of just integrating with cell types within species. We further used the cosine distances for graph visualization of the integration results where distance scores are converted as edge weights.

### Analysis of scRNA-seq Data

Raw sequencing data from cDNA and multiplexed libraries was demultiplexed and aligned with the 10X Genomics Cell Ranger pipeline (v.7.2.0). For mouse experiments, data was aligned to a custom GRCm38/mm10 reference genome including the *tdTomato* transgene used in this study. For human PDX experiments, data was aligned to a combined human (GRCh38) 2020-A and GRCm38/mm10 reference genome.

### Mice

Genetically engineered mouse strains used in this study were maintained in a C57BL/6 x Sv129 mixed background and include the following genetic alleles: *Kras^LSL-G^*^12*D/*+^ [65], *Trp*53*^flox/flox^* [66], and *Rosa*26*^tdTomato/^*^+^ [67]. All mice were maintained monitored by the investigators and veterinary staff at the Research Animal Resource Center at MSKCC with food and water provided ad libitum.

### Patient-derived xenografts (PDXs)

Histologic human UPS samples from MSKCC were obtained under MSKCC IRB #02-060, #06-107, and IRB #12-245. MSK-IMPACT profiling [68] was previously performed. Primary tumors for generation of PDX models were obtained with informed consent from patients under protocols approved by the MSKCC Institutional Review Board as above as well as MSKCC IRB #14-091.

### Autochthonous and Patient-derived xenograft (PDX) models of undifferentiated pleomorphic sarcomas (UPS)

Autochthonous tumors were induced in *Kras^LSL-G^*^12*D/*+^; *Trp*53*^flox/flox^*; and *Rosa*26*^tdTomato/^*^+^ (KPT) mice with 9×107 plaque-forming units (PFU) of AdCMV-Cre (Iowa Viral Vector Core) as previously described [6], in mice that were between 9-24 weeks of age. Immunocompromised NOD.Cg-*Prkdc^scid^ Il*2*rg^tm^*^1*Wjl*^*/SzJ* (aka NSG mice) [69] (The Jackson Laboratory, catalog #005557) mice were used for transplantation of patient-derived xenografts (PDXs). Approximately equal numbers of male and female mice were included in all experimental groups in all mouse experiments where feasible. Mice were treated in accordance to all relevant institutional and national guidelines and regulations, and mice were euthanized by CO2 asphyxiation, followed by intracardiac perfusion with S-MEM (Gibco, catalog #11380-037) to clear tissues of blood when appropriate. All animal studies were approved by the Memorial Sloan Kettering Cancer Center (MSKCC) Institutional Animal Care and Use Committee (protocol # 17-11-008).

### Chemotherapy treatments

Mice were treated with intraperitoneal Doxorubicin (0.9-1 mg/kg) twice a week for up to 4 weeks. NSG mice bearing UPS2236 PDXs were harvested at 3 weeks, approximately 1 week earlier than the preplanned endpoint of 4 weeks due to tumor growth constraints.

### Dissociation of UPS tumors

For isolation of tumor cells from KPT mice and PDXs, mice were euthanized at the indicated time points post tumor induction. Dissected tumors were dissociated with a mixture of Dispase II (Corning, #354235, 0.6 U/ml for GEMMs and 1.2U/ml for PDXs), Collagenase Type IV (Thermo Fisher Scientific, #17104019; 0.167 U/ml for GEMMs and 0.333 U/ml for PDXs), and DNase I (Stemcell Technologies, #07469; 10 U/ml for GEMMs and 20 U/ml for PDXs) in S-MEM solution at 37 *^°^*C as previously described [38]for 1 hr. The dissociated cells were filtered using a 100 *µm* filter and spun at 1000 rpm for 10 min at 4 *^°^*C. The supernatant was removed by aspiration and red blood cell lysis was performed using BD Pharm Lyse (BD Biosciences, #555899) for 1 minute on ice. Cells were then washed with sterile media containing 2% heat inactivated FBS, passed through a 40 *µm* filter, and pelleted at 300 g for 5 min at 4 *^°^*C. The supernatant was removed and cells were resuspended in Fluorescence-Activated Cell Sorting (FACS) buffer media (2% heat-inactivated FBS in PBS) and counted for use in FACS below.

### Flow cytometry and Fluorescence-activated cell sorting (FACS)

Cells were prepared as above and Fc block (BD Biosciences, #553142) was added on ice for 5 min prior to being stained with the appropriate antibody panel (Extended Data Table 1). Cells were incubated for 20 min before being washed with FACS buffer media and pelleted with a 5 min, 300 g spin at 4 *^°^*C. Cells were washed twice and DAPI was added to each sample to identify dead cells. Cell sorting was performed at the Flow Cytometry Core Facility at Sloan Kettering Institute/MSKCC, using a BD FACS Aria Sorter. Cells were sorted using the ‘4-way purity’ mode. Cancer cells from the GEMM model were sorted with (CD45/CD31/CD11b/CD11c/F4/80/TER-119)-/DAPI-(live+)/*tdTomato*^+^ (tumor) gates and cancer cells from the PDX model were sorted with (CD45-Human/CD45-Mouse/TER-119/CD31-Mouse/H-2kd-Mouse)-/DAPI-gates. Cells were sorted into SMEM with 2% heat-inactivated FBS.

### Tissue processing and immunofluorescence staining

Mice were euthanized by CO2 asphyxiation. Tissues were fixed in 10% neutral buffered formalin for 24-48 h at 4 *^°^*C and embedded in paraffin. Immunofluorescence imaging was performed on 5 *µm* formalin-fixed paraffin-embedded (FFPE) sections and sections were stained using a Leica automated stainer. Mounted slides were imaged using the Zeiss Axio Imager Z2 and ZEN 2.3 software or digitally scanned using Mirax Midi-Scanner (Carl Zeiss AG). Image analysis was performed using Fiji software. Due to noted batch effect from multiple rounds of staining of MFH9 samples, a batch correction was performed where chemotherapy treated samples stained for SLC2A1/GLUT1^+^ tumor cells were normalized by the average SLC2A1/GLUT1^+^ signal from the vehicle treated samples in the same batch. Additionally, as necrosis is known to cause nonspecific binding of antibodies, quantification of SLC2A1/GLUT1 signaling was normalized by the percentage of live cells as measured by flow cytometry. Antibodies and dilutions used are available in **Extended Data Table 1**.

### Droplet based scRNA-seq

Single cell suspensions from UPS tumors were prepared and stained as above. Live sorted cells were collected by flow cytometry, washed once with PBS containing 2% heat inactivated FBS (HI-FBS) and resuspended to a final concentration of 700–1,300 cells per *µl* of SMEM + 2% HI-FBS. The viability of cells in all experiments was confirmed to be above 80% by 0.2% (w/v) Trypan Blue staining (Invitrogen, Countess II) and processed by droplet based scRNA-seq using the 10X genomics Chromium Single Cell 3’ Library & Gel bead Kit >=V3 according to manufacturer’s protocol. Following reverse transcription and cell barcoding in droplets, emulsions were broken, and cDNA purified using Dynabeads MyOne SILANE followed by PCR amplification as per manufacturer’s instructions. Between 10,000 to 30,000 cells were targeted for each droplet formulation lane using Chromium. Samples were multiplexed using the TotalSeq B cell hashing protocol [70] (Biolegend, **Extended Data Table 1**). Final libraries were sequenced on Illumina NovaSeq S4 platform (R1 – 28 cycles, i7 – 8 cycles, R2 – 90 cycles).

### Processing and quality control of single cell data

All downstream analysis was performed in Python using the scanpy [71] analysis package and custom scripts. Hashed (i.e. multiplexed) scRNA-seq data from treated and control replicates were separated (using hashsolo[72]), mapped and gene quantified into a combined count matrix. In cases where sample deplutliplexing yielded poor results we used a custom Gaussian mixture model (GMM) approach to define a threshold value for indices background counts from true counts. Cells with >1 uniquely assigned index were removed from subsequent analysis.

Cells with less than 500 Unique molecular identifiers (UMIs), more than 15% mitochondrial UMIs, and low complexity (total counts <30,000 UMI or number of detected genes < 6000) were removed. For PDX analyses, cells with primarily mouse RNA expression were removed. The gene *MALAT1/Malat1* was also removed due to its high ubiquitous expression in all single cell RNA-seq datasets.

Datasets were normalized with scanpy function normalize_total with target_sum=1e4, and then logp1 transformed. Cancer cells were identified by removal of cell populations expressing immune markers (*PTPRC*) and fibroblasts (*DCN*). For studies using mouse models, identification of relevant cell clusters was further refined by detecting the expression of cell-line specific transgene (*tdTomato*).

### Feature selection, dimensionality reduction, clustering and embedding

Highly variable features were selected using a variance stabilizing transformation[73], and dimensionality reduction was performed on normalized, log-transformed count data using principal component analysis. Nearest neighbor networks were calculated on the 50 principal components and clusters based on Leiden clustering and then visualized with Uniform Manifold Approximation and Projection (UMAP).

### Gene signature scoring

For scoring of gene signatures in the PDAC data, classical and basal signatures were compiled from Raghavan and colleagues [32] and mesenchymal signature was collected from HALLMARK_EPITHELIAL_TO_MESENCHYMAL [34]. For scoring of gene signatures in the LUAD data, AT2-like, AT1-like, HPCS, Advanced, EMT, and Endoderm-like signatures were compiled from Marjanovic, Hofree, Chan, etal. [38]. The gene lists were then used in the scanpy.score_genes function to label the cells.

### Benchmarking with other integration algorithms

ScVital was compared to four other algorithms commonly used for scRNA batch correction and cross-species integration: Batch balanced K nearest neighbors (BBKNN) (v1.6.0), Harmony (v0.0.9), scVI (v1.1.2), and scDREAMER. BBKNN is a graph-based integration method that corrects batch effects by increasing the neighborhood size of each cell to include multiple batches and penalize neighborhoods of one batch. This allows BBKNN to merge neighborhoods of similar cell types [11]. Harmony uses an expectation maximization approach on principal components of the data. First, Harmony performs a soft clustering of the data then iteratively merges similar clusters penalizing groups containing only one batch [12]. scVI uses variational autoencoders to integrate the data and model library size. scVI embeds the datasets into the same shared latent space [13]. These methods have been benchmarked [22] and Harmony is shown to work very well on “simple” and scVI is recommended for larger datasets with more “complex” batch effects. Another similar algorithm to scVital is scDREAMER [14] that is similarly based on a variational autoencoder but has 2 adversarially trained networks that train the autoencoder and help remove batch effect from the latent space. Although these algorithms are well suited for overcoming batch effects none are specifically designed for cross-species integration.

### Data Availability

ScRNA-seq raw and processed data reported in this article is available at GEO_XXXX (will be freely available at time of publication). Source code and custom scripts for downloading and running scVital are publicly available under open software license in https://github.com/j-rub/scVital and using pypi. Source code and custom scripts for reproducing the figures in this manuscript and external data used for benchmarking are publicly available at https://github.com/betelab/Rub_scVital

## Author Contributions

**J. Rub:** Conceptualization, Data curation, Formal Analysis, Investigation, Methodology, Software, Validation, Visualization, Writing – original draft; **J.E. Chan:** Conceptualization, Methodology, Investigation, Supervision, Funding acquisition, Validation, Writing – review & editing; **C. Sussman:** Data curation, Investigation; **W.D. Tap:** Supervision, Funding acquisition; **S. Singer:** Resources; **T. Tammela:** Conceptualization, Methodology, Project administration, Resources, Supervision, Validation, Funding acquisition, Writing – review & editing; **D. Betel:** Conceptualization, Methodology, Project administration, Resources, Supervision, Validation, Funding acquisition, Writing – review & editing

## Acknowledgments

This work was supported by NIH R01-AG054720 (to D. Betel), K08-CA267072, the NIH Loan Repayment Program, and the Linn Fund for Sarcoma Research (to J.E. Chan), R01-CA290400 (to T. Tammela), and the NIH/NCI Cancer Center Support Grant P30-CA08748 (to MSKCC). We thank members of the Tammela laboratory for helpful discussions. We would also like to thank Dr. C. Antonescu for her expertise in histological classification of sarcoma. We thank W. Kang, M. Tipping, and the Molecular Cytology Core for histology support; E. Chan and E. Rosiek for help with image analysis and quantification; E. de Stanchina and the Antitumor Assessment Core Facility for support with drug administration and tumor transplant experiments; R. Gardner, M. Kweens, and A. Longhini for FACS support; R. Chaligne and the Single Cell Analytics Innovation Lab for single cell sequencing support, and N. Mohibullah and the Integrated Genomics Operation for next-generation sequencing support. Huge thank you to Doron Haviv for helpful discussions about deep learning. We also want to acknowledge Microsoft copilot in helping to code and comment code.

