## Supplemental Figure 1, 2, 3, 4, 5, 6, 7, 8 for "A deep-learning tool for species-agnostic integration of cancer cell states"

[1]Tri-I Program in Computational Biology & Medicine, Weill Cornell Medicine, New York, NY, 10065, USA.

[2]Cancer Biology and Genetics Program, Sloan Kettering Institute, Memorial Sloan Kettering Cancer Center, New York, NY, 10065, USA.

[3]Division of Solid Tumor Oncology, Department of Medicine, Memorial Sloan Kettering Cancer Center, New York, NY, 10065, USA.

[4]Department of Surgery, Memorial Sloan Kettering Cancer Center and Weill Cornell Medical College, New York, NY, 10065, USA.

[5]Division of Hematology & Medical Oncology, Department of Medicine, Weill Cornell Medicine, New York, NY, 10065, USA.

[6]Institute for Computational Biomedicine, Weill Cornell Medicine, New York, NY, 10065, USA.

December 20, 2024

### Supplementary Figures

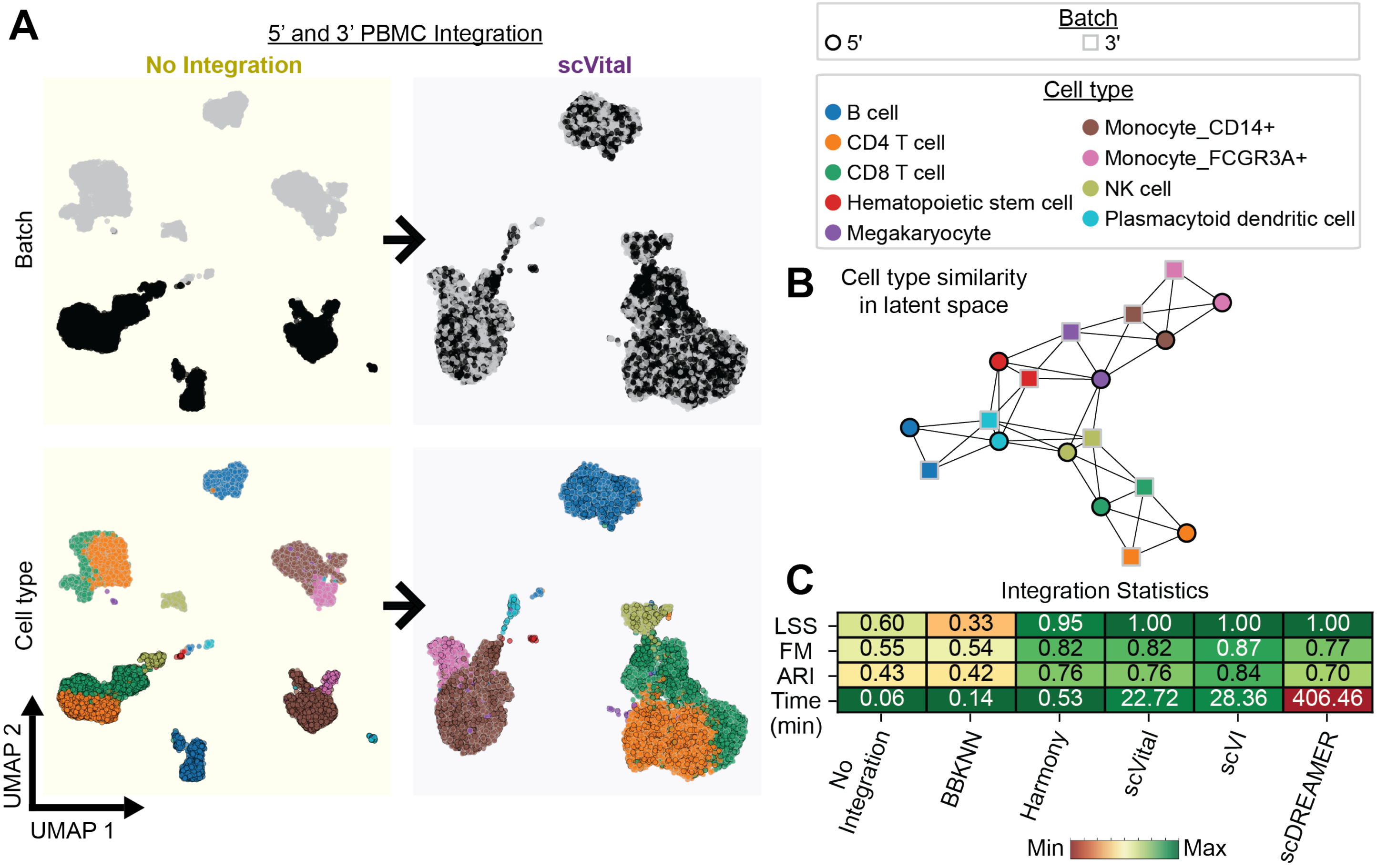


Figure S1: **ScVital Integration of 5’ and 3’ PBMC scRNA-seq data**. **A.** Four UMAP visualizations of 5’ and 3’ PBMC data with no integration in yellow (left) and scVital integration in purple (right) colored by species (top) and cell type (bottom). Nodes in the UMAP represent cells where their outline represents the species (black for 5’ and gray for 3’) and their internal color represents the cell type. With no integration there is a clear separation of PBMC data based on sequencing direction with no overlap of homologous cell types. After integration with scVital there is overlap of the same cell types as shown by the same internal color overlapping in the UMAP visualization. **B.**Graph visualization of LSS. Each node represents the average latent space of a cell type after scVital integration where the outline color is the batch and internal color is the true label as determined by the input dataset. The distances of the nodes represent the cosine similarities of the cell type’s average latent space compared to other cell types. Same-colored nodes with black and gray outlines indicates integration of batch and conservation of cell type. **C.** Integration Statistics. LSS, FM, ARI and runtime (in minutes) of 5’ and 3’ PBMC data integrated using no integration, BBKNN, Harmony, scVital, scVI, and scDREAMER. ARI and FM metrics are the lowest for No Integration. ScVital performs similar to the other gold standard integration algorithms achieving similar ARI, FM, and LSS scores.

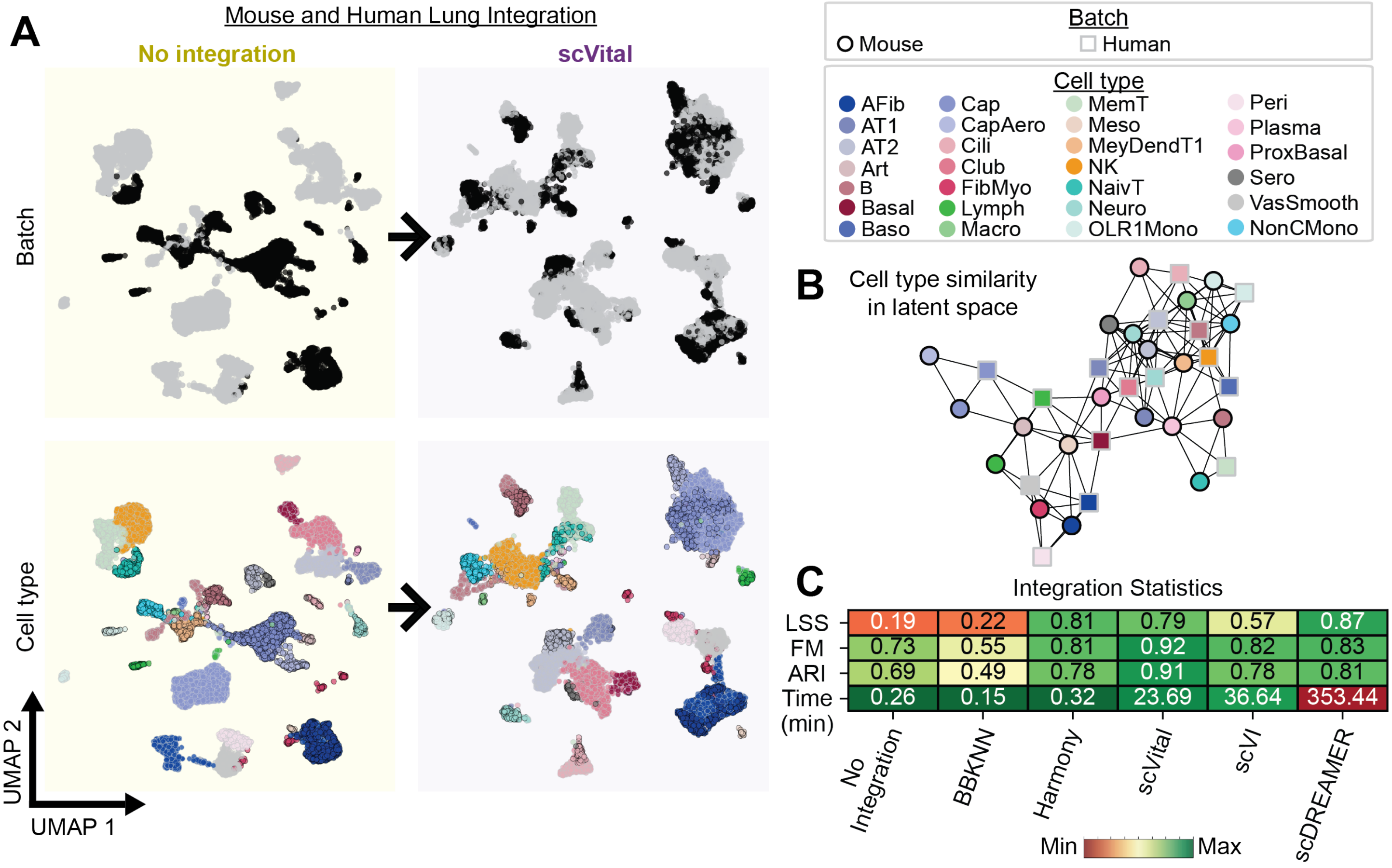


Figure S2: **ScVital Integration of Normal Mouse and Human Lung scRNA-seq data. A.** Four UMAP visualizations of mouse and human lung data with no integration in yellow (left) and scVital integration in purple (right) colored by species (top) and cell type (bottom). Nodes in the UMAP represent cells where their outline represents the species (black for mouse and gray for human) and their internal color represents the cell type. With no integration there is a clear separation of mouse and human lung data with no overlap of homologous cell types. After integration with scVital there is overlap of similar cell types as shown by the same internal color overlapping in the UMAP visualization. Alveolar fibroblast(AFib), artery(Art), Alveolar type 2(AT2), proliferating basal(Baso), proximal basal(ProxBasal), general capillary cell(Cap), capillary aerocyte(CapAero), ciliated(Cili), fibromyocyte(FibMyo), lymphatic(Lymph), memory T cell(MemT), mesothelial(Meso), Dendrocyte(MeyDendT1), naive T cell (naivT), neuroendocrine(Neuro), OLR1+ monocyte(OLR1Mono), pericyte(Peri), serous(Sero), vascular smooth muscle(VasSmooth), non-classical Mmnocyte(NonCMono). **B.** Graph visualization of LSS. Each node represents the average latent space of a cell type after scVital integration where the outline color is the batch and internal color is the true label as determined by the input dataset. The distances of the nodes represent the cosine similarities of the cell type's average latent space compared to other cell types. Same-colored nodes with black and gray outlines indicates integration of species and conservation of cell type. **C.** Integration Statistics. LSS, FM, ARI and runtime (in minutes) of mouse and human lung integrated using no integration, BBKNN, Harmony, scVital, scVI, and scDREAMER. No integration yields the lowest integration metrics. ScVital performs comparably well to the other gold standard integration algorithms achieving similar ARI, FM, and LSS scores.

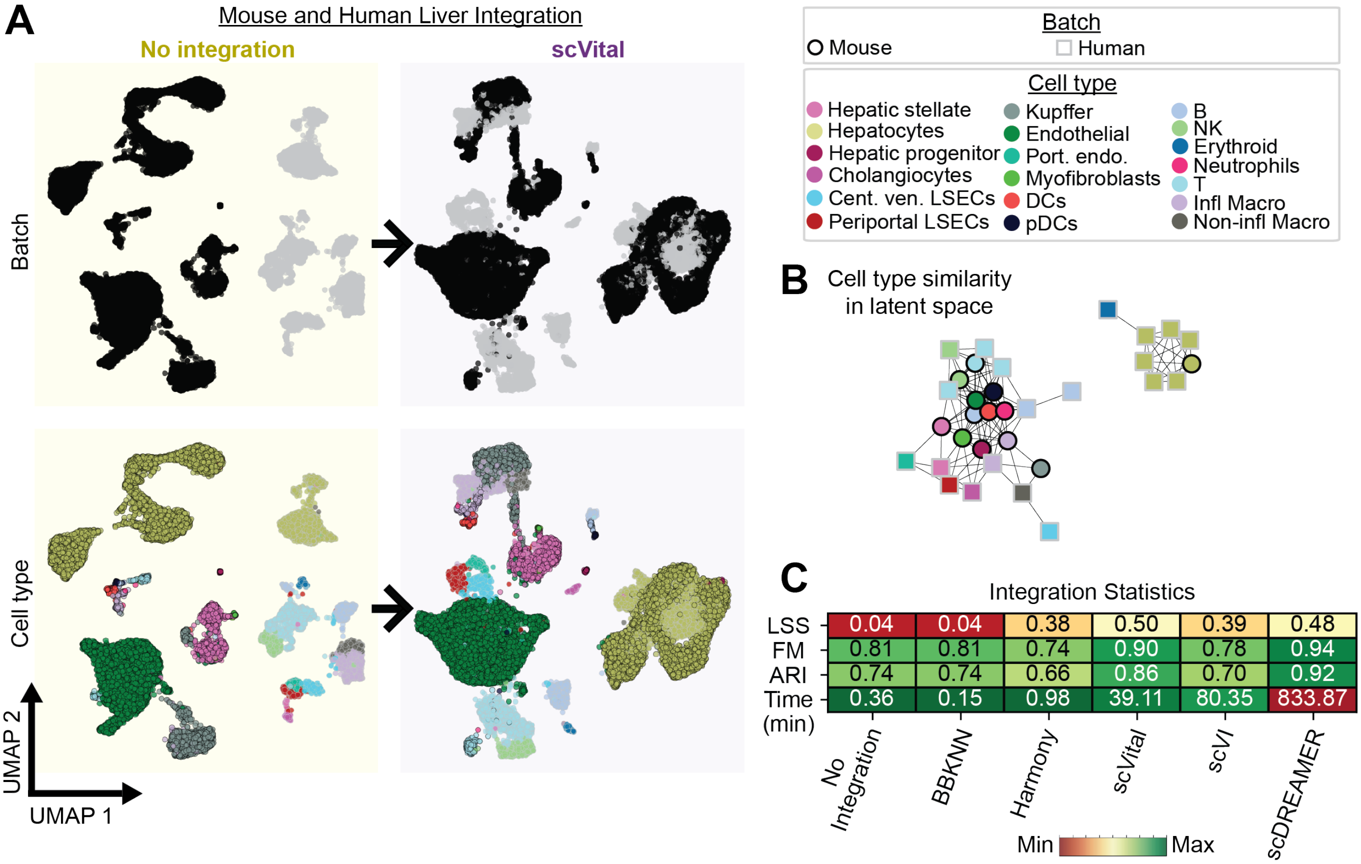


Figure S3: **ScVital Integration of Normal Mouse and Human Liver scRNA-seq data. A.** Four UMAP visualizations of mouse and human liver data with no integration in yellow (left) and scVital integration in purple (right) colored by species (top) and cell type (bottom). Nodes in the UMAP represent cells where their outline represents the species (black for mouse and gray for human) and their internal color represents the cell type. With no integration there is a clear separation of mouse and human liver data with no overlap of homologous cell types. After integration with scVital there is overlap of similar cell types as shown by the same internal color overlapping in the UMAP visualization. Dendritic Cells (DCs), portal endothelial (port. endo.), central venous liver sinusoidal endothelial cell (port. cent. LSEC), and inflamitory macrophages (infl. Macro). **B.** Graph visualization of LSS. Each node represents the average latent space of a cell type after scVital integration where the outline color is the batch and internal color is the true label as determined by the input dataset. The distances of the nodes represent the cosine similarities of the cell type's average latent space compared to other cell types. Same-colored nodes with black and gray outlines indicates integration of species and conservation of cell type. **C.** Integration Statistics. LSS, FM, ARI and runtime (in minutes) of mouse and human liver integrated using no integration, BBKNN, Harmony, scVital, scVI, and scDREAMER. No integration yields the lowest integration metrics. ScVital performs comparably well to the other gold standard integration algorithms achieving similar ARI, FM, and LSS scores.

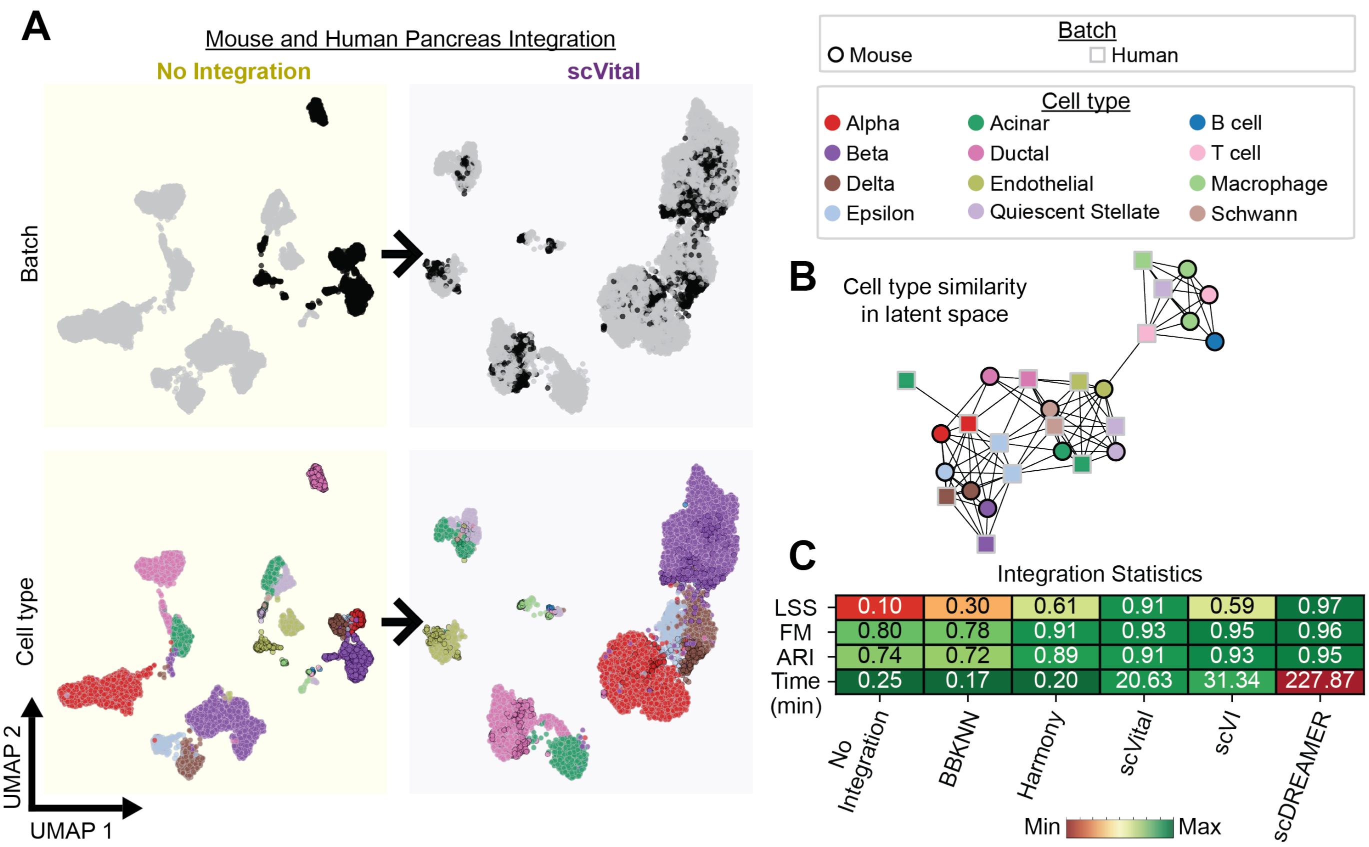


Figure S4: **ScVital Integration of Normal Mouse and Human Pancreas scRNA-seq data A.** Four UMAP visualizations of mouse and human pancreas data with no integration in yellow (left) and scVital integration in purple (right) colored by species (top) and cell type (bottom). Nodes in the UMAP represent cells where their outline represents the species (black for mouse and gray for human) and their internal color represents the cell type. With no integration there is a clear separation of mouse and human pancreas data with no overlap of homologous cell types. After integration with scVital there is overlap of similar cell types as shown by the same internal color overlapping in the UMAP visualization. **B.** Graph visualization of LSS. Each node represents the average latent space of a cell type after scVital integration where the outline color is the batch and internal color is the true label as determined by the input dataset. The distances of the nodes represent the cosine similarities of the cell type's average latent space compared to other cell types. Same-colored nodes with black and gray outlines indicates integration of species and conservation of cell type. **C.** Integration Statistics. LSS, FM, ARI and runtime (in minutes) of mouse and human pancreas integrated using no integration, BBKNN, Harmony, scVital, scVI, and scDREAMER. No integration yields the lowest integration metrics. ScVital performs comparably well to the other gold standard integration algorithms achieving similar ARI, FM, and LSS scores.

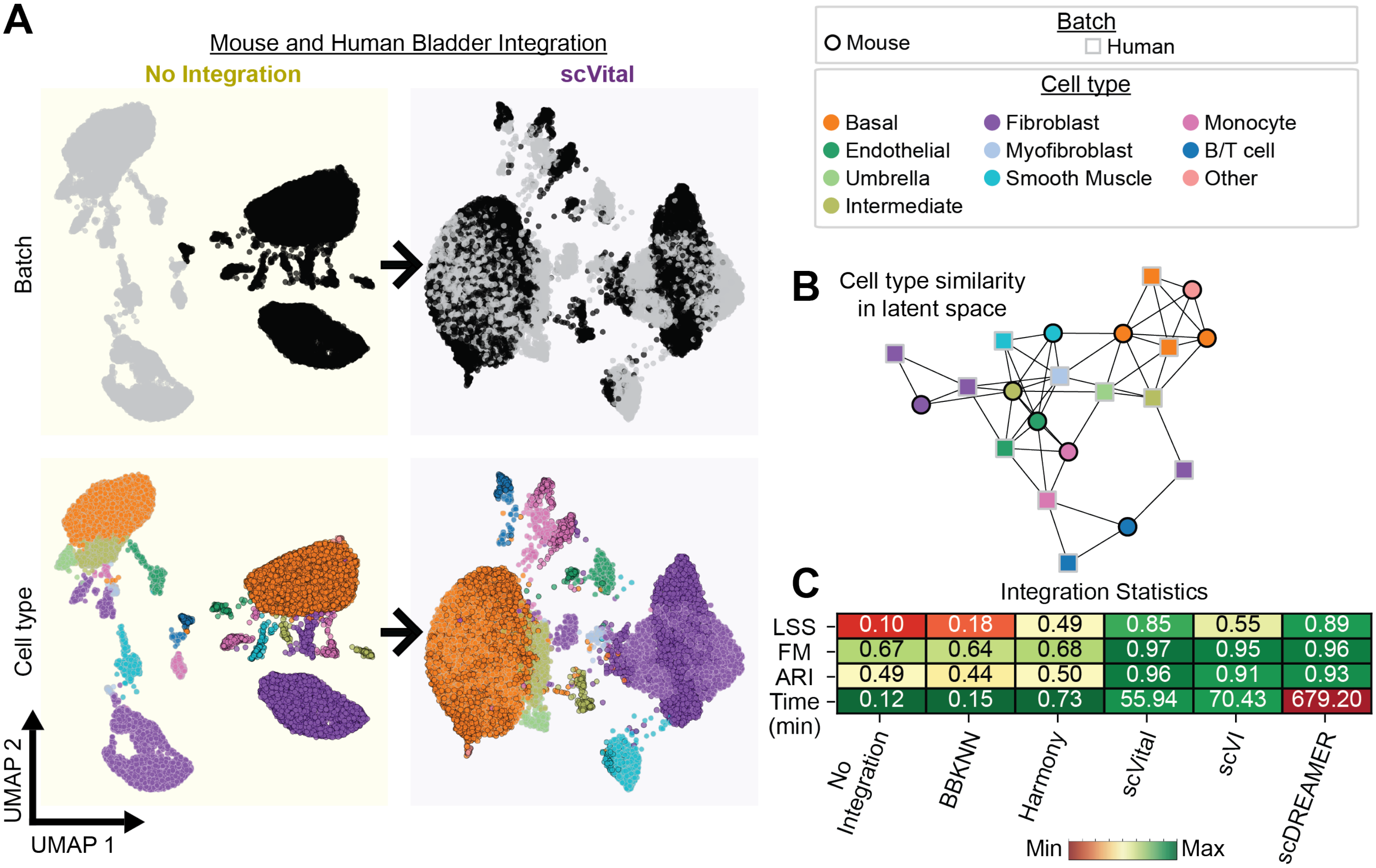


Figure S5: **ScVital Integration of Normal Mouse and Human Bladder scRNA-seq data. A.** Four UMAP visualizations of mouse and human bladder data with no integration in yellow (left) and scVital integration in purple (right) colored by species (top) and cell type (bottom). Nodes in the UMAP represent cells where their outline represents the species (black for mouse and gray for human) and their internal color represents the cell type. With no integration there is a clear separation of mouse and human bladder data with no overlap of homologous cell types. After integration with scVital there is overlap of similar cell types as shown by the same internal color overlapping in the UMAP visualization. **B.** Graph visualization of LSS. Each node represents the average latent space of a cell type after scVital integration where the outline color is the batch and internal color is the true label as determined by the input dataset. The distances of the nodes represent the cosine similarities of the cell type's average latent space compared to other cell types. Same-colored nodes with black and gray outlines indicates integration of species and conservation of cell type. **C**. Integration Statistics. LSS, FM, ARI and runtime (in minutes) of mouse and human bladder integrated using no integration, BBKNN, Harmony, scVital, scVI, and scDREAMER. No integration yields the lowest integration metrics. ScVital performs comparably well to the other gold standard integration algorithms achieving similar ARI, FM, and LSS scores.

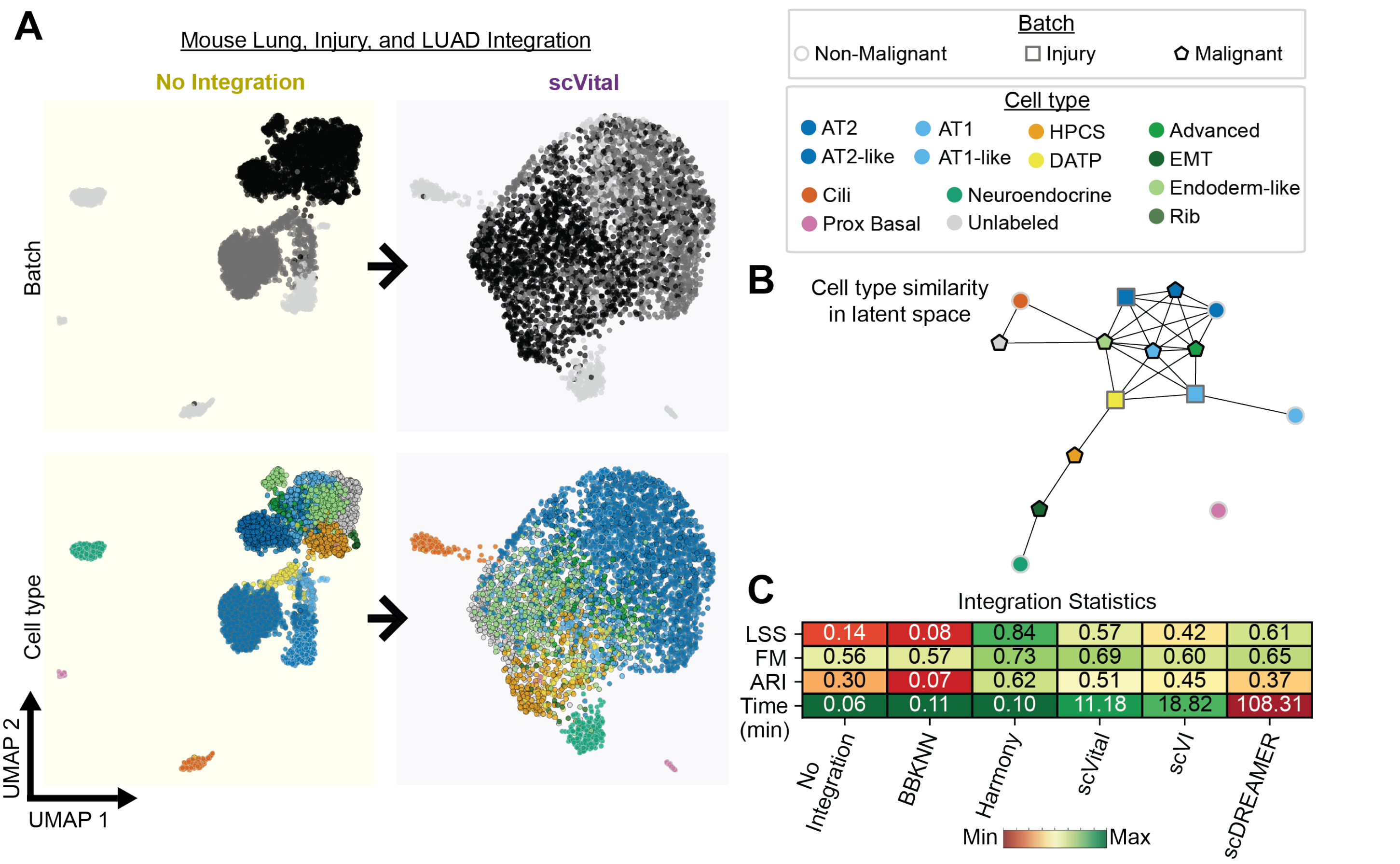


Figure S6: **ScVital Integration of Mouse normal Lung, injured lung, and LUAD scRNA-seq data.** **A.** Four UMAP visualizations of mouse normal Lung, injured lung, and LUAD data with no integration in yellow (left) and scVital integration in purple (right) colored by species (top) and cell type (bottom). Nodes in the UMAP represent cells where their outline represents the species (light gray for mouse, dark gray for injury, and black for LUAD) and their internal color represents the cell type. With no integration there is a clear separation of mouse normal Lung, injured lung, and LUAD data with no overlap of homologous cell types. After integration with scVital there is overlap of similar cell types as shown by the same internal color overlapping in the UMAP visualization. **B.** Graph visualization of LSS. Each node represents the average latent space of a cell type after scVital integration where the outline color is the batch and internal color is the true label as determined by the input dataset. The distances of the nodes represent the cosine similarities of the cell type's average latent space compared to other cell types. Same-colored nodes with black and gray outlines indicates integration of species and conservation of cell type. **C.** Integration Statistics. LSS, FM, ARI and runtime (in minutes) of mouse normal Lung, injured lung, and LUAD integrated using no integration, BBKNN, Harmony, scVital, scVI, and scDREAMER. No integration yields the lowest integration metrics. ScVital performs comparably well to the other gold standard integration algorithms achieving similar ARI, FM, and LSS scores.

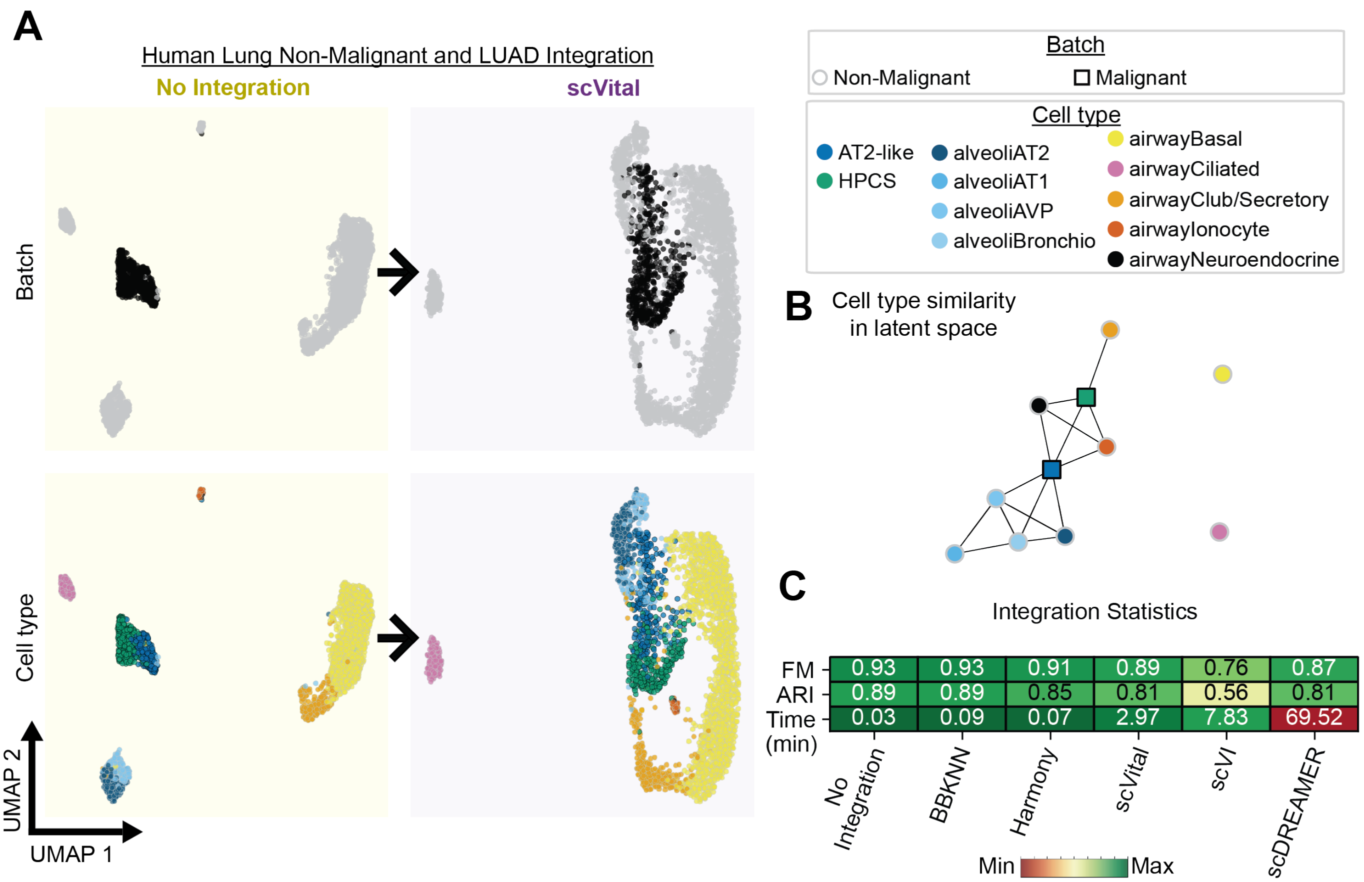


Figure S7: **ScVital Integration of Human normal Lung and LUAD scRNA-seq data. A.** Four UMAP visualizations of human normal Lung and LUAD data with no integration in yellow (left) and scVital integration in purple (right) colored by species (top) and cell type (bottom). Nodes in the UMAP represent cells where their outline represents the species (black for malignant and gray for non-malignant) and their internal color represents the cell type. With no integration there is a clear separation of human normal lung and LUAD data with no overlap of homologous cell types. After integration with scVital there is overlap of similar cell types as shown by the same internal color overlapping in the UMAP visualization. **B.** Graph visualization of LSS. Each node represents the average latent space of a cell type after scVital integration where the outline color is the batch and internal color is the true label as determined by the input dataset. The distances of the nodes represent the cosine similarities of the cell type's average latent space compared to other cell types. Same-colored nodes with black and gray outlines indicates integration of species and conservation of cell type. **C.** Integration Statistics. LSS, FM, ARI and runtime (in minutes) of human normal lung and LUAD integrated using no integration, BBKNN, Harmony, scVital, scVI, and scDREAMER. No integration yields the lowest integration metrics. ScVital performs comparably well to the other gold standard integration algorithms achieving similar ARI, FM, and LSS scores.

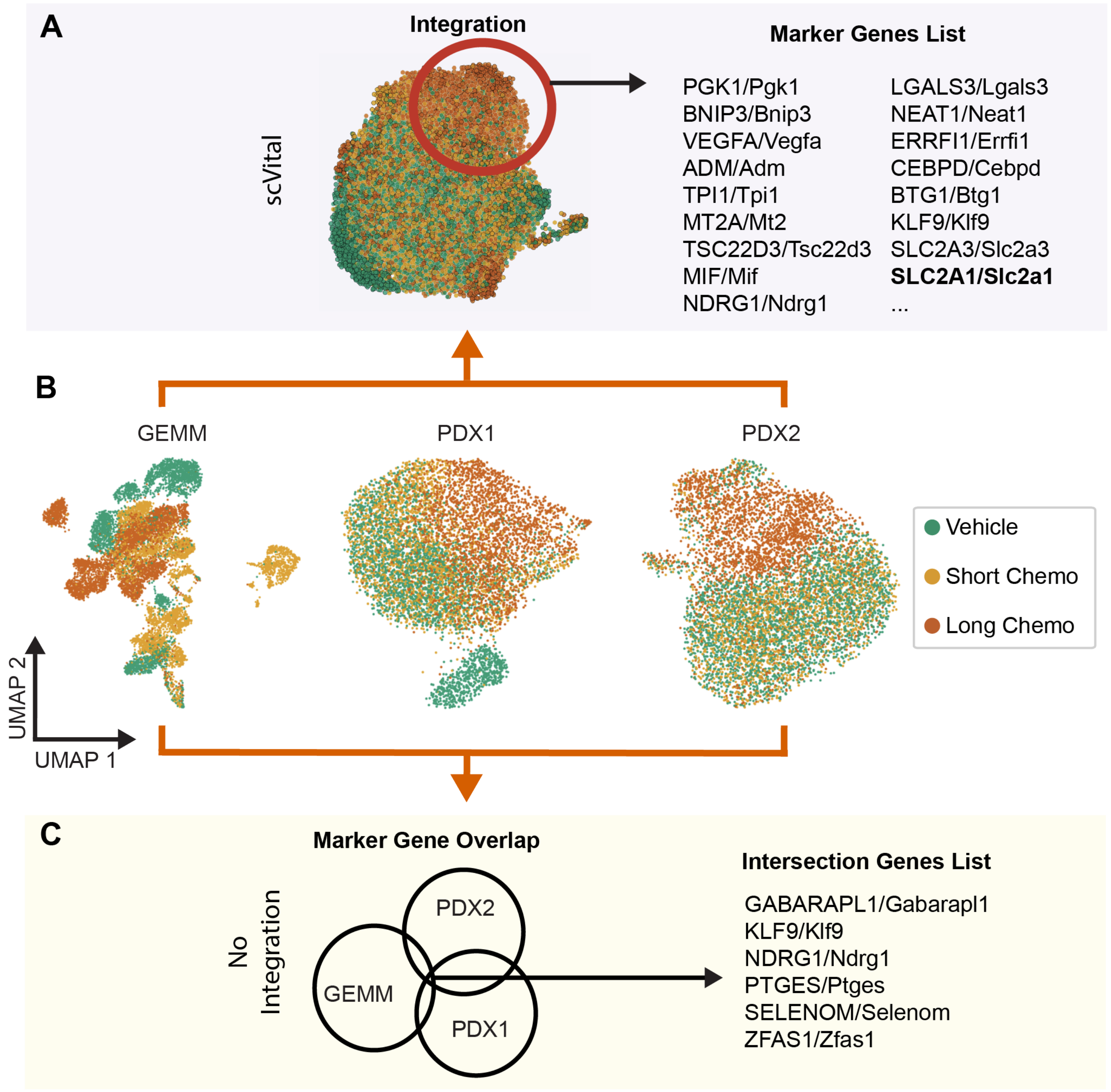


Figure S8: **Determining overlapping genes for UPS GEMM and 2 PDXs with scVital and with the normal workflow.** **A.** Integration of GEMM and 2 PDX of UPS with scVital identifies a substantial gene list for an overlapping cluster dominated by long chemotherapy treatment circled in red including *SLC2A1/Slc2a1*, which is a gene marker for hypoxia. **B.** Individual UMAP visualization of GEMM and 2 PDXs of UPS colored by treatment strategy. **C.** During the standard workflow of calculating overlapping genes of the long-term chemotherapy treatment groups identify 6 genes that are in common with the 3 groups. These genes do not have an obvious relationship.
